# Lipid Droplets of Human Carotid Atherosclerotic Plaques: Cholesteryl Arachidonate Core and Cytoskeletal Fibrotic Cover

**DOI:** 10.1101/2025.08.18.670831

**Authors:** Yilu Xu, Yang Wang, Guangxin Yang, Qichen Feng, Xiaochuan Fu, Qiumin Liao, Xiao-Juan Wei, Xia Chen, Hongchao Zhang, Shuyan Zhang, Changming Wang, Pingsheng Liu

**Author notes:** These authors contributed equally (Yilu Xu, Yang Wang, Guangxin Yang, Qichen Feng).

## Abstract

Lipid droplets (LDs) within atherosclerotic plaques are central to cardiovascular disease, yet their molecular composition remains largely unknown. More than 20 years experiences of developing LD purification method allowed us recently to successfully purify LDs from atherosclerotic plaques of patients undergoing carotid artery surgery, including three types, soft (lipid), mixed, and hard (calcified). In addition, the subpopulations of large LDs and small LDs were purified and studied, respectively. We report the first comprehensive proteo-lipidomic analysis of these LDs. Unexpectedly, about 19.3% of the plaque LD proteins belonged to cytoskeleton, while the content of lipid metabolism enzymes on plaque LDs was low. Our results show that the compositions of lipids and proteins were similar between two population LDs. Compared to soft (lipid) plaque LDs, LDs from mixed and hard (calcified) plaques had fewer metabolic enzymes and more cytoskeletal proteins. Lipidomics of the LD core uncovered an unexpected enrichment of polyunsaturated fatty acid cholesteryl esters (PUFA-CEs), which constituted 88.3% of all cholesterol esters. Notably, esters of the pro-inflammatory precursor arachidonic acid (AA) were among the most abundant species (16.5%). In addition, plaque LD were uniquely enriched in phospholipids including phosphatidic acid (PA), phosphatidylethanolamine plasmagens (PE-P), sphingomyelin (SM), and phosphatidylserine (PS). Our findings redefine the plaque LD as a dual-function “bad” organelle: a metabolically inert container for cholesterol ester, coated with cytoskeletons, and a massive reservoir for inflammatory precursors.

## Introduction

Atherosclerotic cardiovascular disease (ASCVD) continues to be a leading cause of morbidity and mortality worldwide, and the mortality rates are steadily climbing^[1]^. Recently, the Lancet Commission re-evaluated coronary artery disease and reclassified this group of diseases as atherosclerotic coronary artery disease (ACAD), underscoring the need for a more systematic and comprehensive understanding of atherosclerosis^[1]^. Lipoproteins play an important role in the onset and progression of atherosclerosis. Macrophages engulf modified low-density lipoprotein (LDL) and transform into foam cells, a hallmark of atherosclerotic lesions^[2]^. This process is accompanied by endothelial dysfunction, lipid accumulation, and inflammation^[3–5]^. Therefore, large amounts of lipoproteins and lipids engulfed by macrophages accumulate within the atherosclerotic plaques. These lipids predominantly exist as lipid droplets (LDs), which become exposed to the extracellular matrix when cells in the plaque core undergo necrosis during lesion progression^[6–8]^.

LDs are unique organelles consisting of a neutral lipid core surrounded by a phospholipid monolayer membrane with resident and dynamic proteins. LDs play important roles in lipid storage, metabolism, and transport. LDs are also involved in protein storage and degradation in addition to storing neutral lipids^[9–13]^. The proteomic and lipidomic profiles of LDs from various tissues and cell types have been extensively investigated following the establishment of reliable isolation methods^[14–16]^, thereby advancing our understanding of LD functions and their roles in biological processes and metabolic diseases. However, LDs from atherosclerotic plaques were first isolated by LANG *et al*. in 1970, and due to technical limitations, only their lipid composition were analyzed, while their protein composition/proteome and detailed lipidome remained unknown^[17]^. With the development of mass spectrometry (MS) technology, analyses of both protein and lipid composition in whole atherosclerotic plaques have been increasingly reported^[18–20]^. Nevertheless, the detailed and comprehensive proteomic and lipidomic characterization of plaque-derived LDs has yet to be performed.

Since 2003 our group has expended major effort on development and improvement of LD isolation and purification methods. We have modified our original method to isolate/purify and omics studied most cell types and tissue samples, including skeletal muscles, liver, and adrenal from human and mouse, monkey skeletal muscle, heart, brown adipose tissue, and liver, mouse brown adipose tissue and testis, rat heart, fish liver, etc. These improved technologies and experiences allowed us to successfully isolate LDs from distinct types of human atherosclerotic plaques and conduct the first in-depth, quantitative proteomic and lipidomic analysis. Our investigation reveals that plaque LDs are unique, fibrotic organelles, with a distinct proteome and a highly unusual, pro-inflammatory lipid signature. These findings provide an unprecedented view into the role of LDs in atherosclerosis and identify novel molecular features that drive plaque pathology.

## Materials and methods

### Materials

PageRuler Prestained Protein Ladder were from ThermoFisher SCIENTIFIC. Polyvinylidene fluoride (PVDF) membranes were obtained from Millipore Life Sciences. Glutaraldehyde, osmium tetraoxide, uranyl acetate and lead citrate were purchased from Electron Microscopy Sciences. Antibodies against GAPDH, Calnexin, β-actin, NCEH1, ADRP, PLIN3, Annexin A2, β-tubulin and ACSL3 were from ABclonal. Antibodies against ATGL was from Cell Signaling Technology. Antibody against Keratin1, vimentin, and α-actin were from Santa Cruz Biotechnology. Antibody against VDAC was from Millipore.

### Human carotid endarterectomy samples

The study was approved by the ethics committee of and was conducted with patient consent and all methods were performed according to approved ethical guidelines. Clinical characteristics (including routine blood tests and liver function tests) and imaging results from Peking University Third Hospital. All 19 plaque samples were obtained from patients who underwent carotid endarterectomy (CEA) at Peking University Third Hospital.

### Histology

Human plaque samples were fixed with 4% paraformaldehyde, embedded in paraffin or frozen, and then sectioned. The frozen sections stained with Oil Red O for visualizing LDs. The paraffin blocks were then cut into 5 µm thick sections and stained with hematoxylin and eosin (H&E). Images were captured with Leica Aperio Versa 200 microscope (Leica, Germany). In addition, paraffin sections were incubated with 3% H_2_O_2_ for 10 min to eliminate endogenous peroxidase activity, and antigen retrieval was performed with 0.3% sodium citrate and phosphate buffer (pH 7.4). Sections were blocked with 1% defatted albumin, immunostained with primary antibodies, and developed after incubation with enzyme-labeled secondary antibodies.

### Isolation of lipid droplets from plaques and extraction of their proteins and lipids

Lipid droplets (LDs) were isolated and analyzed from human plaque using a modified protocol previously described^[15]^. Briefly, the collected plaque tissue was first washed with saline to remove blood, then the endothelial cells and other parts were removed, and the plaques were minced with scissors and scalpels, and homogenized with a Tenbroeck tissue grinder (WHEATON, 357424) in 10 mL Buffer A (20 mM Tricine, pH 7.6, 250 mM Sucrose) containing 0.2 mM PMSF. The homogenate was centrifuged at 2,000*g* for 10 min to obtain large LDs. The above remaining supernatant was centrifuged at 182,000*g* at 4°C for 1 h. The resulting precipitate and intermediate liquid are the total membrane (TM) and cytosolic fractions, respectively. Small LDs formed an upper layer and were carefully collected into a 1.5 mL Eppendorf tube. The collected large and small LDs were washed by centrifugation at 2,000*g* and 21,380*g* for 3 min at 4°C, respectively, and the underlying solution was discarded. LDs were gently resuspended in 200 µL Buffer B (20 mM HEPES, 100 mM KCl, and 2 mM MgCl_2_, pH 7.4). This washing step was repeated three times. The TM fraction was also washed 2–3 times with Buffer B by centrifugation at 21,380*g* for 5 min at 4°C.

Lipid extraction and protein precipitation from LDs were performed using acetone/chloroform (4:1, v/v) treatment followed by centrifugation at 21,380*g* for 10 min at 4°C. The resulting protein pellet was then dissolved in 2× SDS sample buffer [100 mM Tris-HCl (Ph 6.8), 4% SDS (m/v), 20% glycerol (v/v), 4% 2-mercaptoethanol and 0.04% bromophenol blue] or stored in an EP tube filled with nitrogen gas at −20°C.

### Transmission electron microscopy (TEM) analysis

Isolated LDs were loaded onto grid first. Then the grid was placed onto a drop of 2.5% glutaraldehyde solution (0.1 M PB, pH 7.2) for 10 min followed by the addition of a drop of 1% osmium tetraoxide solution (0.1 M PB, pH 7.2), which was incubated for 10 min. Then, the LDs were stained with 0.1% tannic acid for 10 min followed by 2% uranyl acetate for 10 min. the grid was washed with water in between each staining step. Then the grids were observed under a Tecnai Spirit electron microscope (FEI, Netherlands).

The collected plaque samples were fixed overnight at 4°C in 2.5% glutaraldehyde (v/v) in (0.1 M PB, pH 7.2) overnight at 4°C. Subsequently, the samples were post-fixed for 2 h at 4°C in 1% osmium tetraoxide with 1.5% potassium ferrocyanide. Following fixation, the samples were dehydrated through an ethanol series and then embedded in Embed 812 to be prepared as 70-nm-thick ultrathin sections. After staining with uranyl acetate and lead citrate, the sections were observed under a Tecnai Spirit electron microscope (FEI, Netherlands).

### Confocal microscopy

Purified plaque LDs were stained with LipidTOX Green for 30 min, and 2 µL of the LD solution was placed on a glass slide, covered with a coverslip, and imaged using an Olympus FV3000 confocal microscope (Olympus Corp, Lake Success, NY).

### Silver staining and Western blotting

For silver staining, proteins from different plaque fractions were dissolved in 2× SDS sample buffer and denatured at 95°C for 5 min. Protein separation is then performed using SDS-polyacrylamide gel electrophoresis (SDS-PAGE). The gel is subsequently fixed in a fixative solution (water: ethanol: acetic acid, 5:4:1, v/v/v) at room temperature for 30 min, followed by sensitization with a sensitizing solution containing 30% ethanol (v/v), 12.7 mM sodium thiosulfate pentahydrate, 0.83 M sodium acetate trihydrate for 30 min. The gels were then washed four times with ddH_2_O, 5 min per wash. After washing, the gels were incubated in silver staining solution (14.72 mM silver nitrate with 4.93 µM formaldehyde added immediately before use) for 20 min. This was followed by development in chromogenic solution (0.24 M sodium carbonate with 4.93 µM formaldehyde added immediately before use) until the protein bands are visible. The reaction was terminated by adding stop solution containing 43.43 mM EDTA-Na_2_ once the bands were clearly developed.

For Western blotting, proteins of different fractions were separated by SDS-PAGE and transferred to 0.2 µm PVDF membrane. The membrane was blocked with 5% BSA for 1 h at room temperature. The primary antibody is incubated with the membrane at room temperature for 1 h or at 4°C overnight. The membrane was then washed three times with washing buffer, each wash lasting 5 min. The secondary antibody was incubated with the membrane at room temperature for 1 h. Following this, the membrane was washed three times with washing buffer for 5 min each and then detected by enhanced chemiluminescence (ECL).

### *In vitro* recruitment assay for plaque LDs

Purified plaque LDs were aliquoted for different treatments. For cytosol incubation, LDs were incubated with cytosol (with or without 1 mM CaCl_2_ and 5 mM EGTA) at 37°C for 60 min, then re-isolated and proteins were extracted with chloroform:acetone (200 µL:800 µL) for analysis by silver staining and Western blotting with the indicated antibodies. For trypsin digestion, LDs were incubated with trypsin at the concentration of 0.025% (w/v) at 37°C for 30 min, then re-isolated and proteins were extracted for analysis. LDs from normal mouse liver were used as controls.

### Thin-layer chromatography (TLC)

After the lipids from the plaque LDs were extracted using acetone/chloroform (4:1, v/v), they were dried under a nitrogen stream. The dried lipid extract was then re-dissolved in chloroform and spotted onto a silica gel thin-layer chromatography (TLC) plate. Neutral lipids were separated using a solvent system of hexane/diethyl ether/acetic acid (80:20:1, v/v/v) and visualized by exposure to iodine vapor.

### Liquid chromatography-tandem mass spectrometry (LC-MS/MS)

Lipids from plaque LDs were extracted and proteins were precipitated according to the method described above. Nitrogen gas was added to the EP tubes containing proteins and stored at −20°C for proteomics analysis, and lipids were dried under a nitrogen stream and stored at −20°C for lipidomics analysis. For proteomics, the LD protein pellet and WTL was dissolved in 20 µL of freshly prepared 8 M urea or was dissolved in 2× sample buffer. For identification of WTL proteins and major protein bands in LDs, proteins dissolved in 2× sample buffer were separated by SDS-PAG and visualized by silver staining. The indicated bands or entire lane were sliced out, reduced with 200 mM dithiothreitol (DTT) solution and incubating at 37°C for 1 h. For identification of LD proteins, frozen LD protein powder was also reduced with 200 mM dithiothreitol (DTT) solution and incubated at 37°C for 1 h. The different sample was diluted 4 times by adding 25 mM ammonium bicarbonate (ABC) buffer. Then adding trypsin (trypsin: protein = 1:50) and incubating at 37°C overnight. The next day, adding 50 µL 0.1% formic acid (FA) to terminate the digestion. The tryptic peptide mixtures were analyzed by using a quadrupole Orbitrap mass spectrometer (Q Exactive HF-X, Thermo Fisher Scientific, Bremen, Germany) operated in a data-independent acquisition with parallel accumulation and serial fragmentation mode. Quantitative analysis peptides from digestion of the original proteins were identified by comparison with the UniProtKB protein database.

For lipidomics, a mixture containing internal standard lipids was added to the lipid sample of LDs and analyzed using ultra-high performance liquid chromatography (ExionLC™ AD) combined with a triple quadrupole mass spectrometer (QTRAP® 6500+). Each ion pair was scanned and detected based on the optimized declustering potential (DP) and collision energy (Collision Energy). Based on the self-built database MWDB (metware database), qualitative analysis was performed based on the retention time RT (Retention time) of the detected substance and the information of the daughter ion and parent ion pair. Lipid quantification was completed using the multiple reaction monitoring (MRM) analysis of the triple quadrupole mass spectrometer. In the MRM mode, the quadrupole first screens the precursor ions (parent ions) of the target substance, and excludes the ions corresponding to other molecular weight substances to preliminarily eliminate interference; the precursor ions are induced by the collision chamber and then broken to form many fragment ions. The fragment ions are then filtered through the triple quadrupole to select the required characteristic fragment ion, eliminating non-target ion interference, making the quantification more accurate and more repeatable. After obtaining the lipid mass spectrometry data of different samples, the peak areas of all substance chromatographic peaks were integrated and quantitative analysis was performed using the internal standard method.

### Data analysis and statistical treatment

Protein family and pathway analysis was performed using the COG (Clusters of Orthologous Groups) and KEGG (Kyoto Encyclopedia of Genes and Genomes) databases, and enrichment analysis was performed on GO and KEGG using enrichment pipelines. These analyses were performed using the DAVID online platform (https://davidbioinformatics.nih.gov/home.jsp). The statistical analyses were performed using Office Excel 2010 (Microsoft Corp.) and GraphPad Prism 8 (NIH, USA). Determination of significance between groups was performed using unpaired Student t-test. Values presented are means ± SD.

## Results

### Human carotid atherosclerotic plaques contain numerous small lipid droplets

We collected 19 human carotid atherosclerotic plaques from patients who underwent carotid endarterectomy (CEA). The clinical characteristics of these patients are summarized in Table 1. Based on CT angiography, the plaques were classified into lipid-rich (lipid), mixed (mix), and calcified (Ca) types(Figure 1A)^[21]^. HE and Oil Red O staining of plaque samples showed that plaques were rich in lipids (Figure 1B). Transmission electron microscopy (TEM) of the ultrathin sections also revealed abundant LDs within the plaques, most of which were small, with diameters under 500 nm. No intact cellular structures were observed in the plaque core, suggesting that these LDs were “extracellular LDs”. In addition, fibrous structures adjacent to LDs were visible (Figure 1C), consistent with previous reports^[7,22]^.

**Figure 1.**
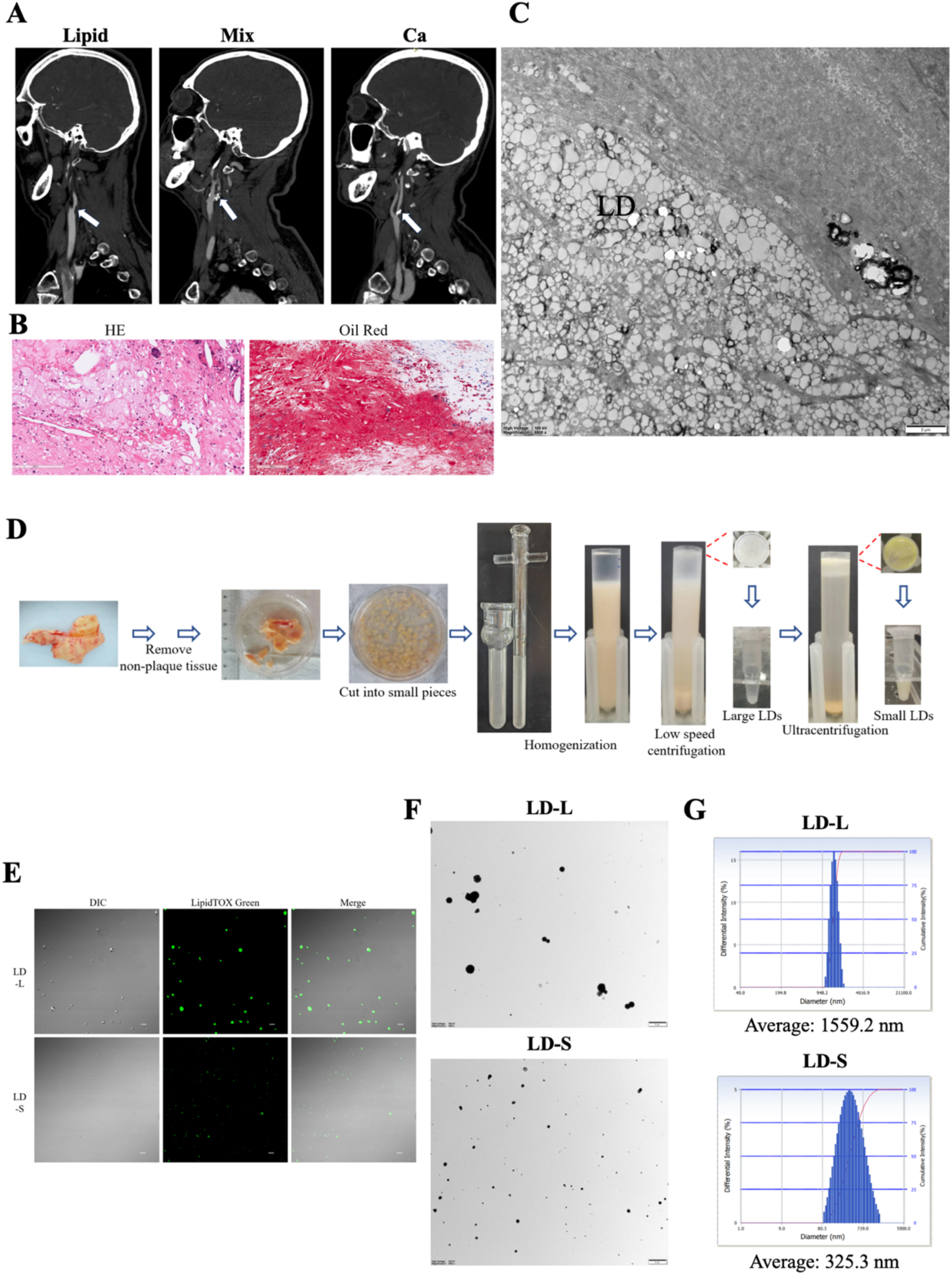
Observation and purification of human carotid atherosclerotic plaque lipid droplets. A. Representative CT angiography of lipid-rich (Lipid), mixed (Mix), and calcified (Ca) human carotid artery plaques. B. Representative H&E and Oil Red O staining of human carotid artery endarterectomy plaque samples. C. Representative TEM images of ultrathin sections of human carotid artery endarterectomy plaque samples. LD, lipid droplets. D. Isolation and purification schematic of plaque LDs. E. Observation of purified plaque LDs by fluorescence microscopy. Purified plaque LDs were stained with LipidTOX green and observed under fluorescence microscope. LD-L: large LDs; LD-S: small LDs. F. TEM observation of positive staining of isolated plaque LDs. G. Size analysis of purified plaque LDs. The isolated LDs were analyzed for size distribution using a Delsa Nano C particle analyzer.

**Table 1.** Basic characteristics of patients with endarterectomies.

| Variable | Plaque classification |  |  |
| --- | --- | --- | --- |
|  | Lipid (n=9) | Mix (n=8) | Ca (n=2) |
| Age (years) | 65±5 | 66±5 | 69/70 |
| Male sex-no. (%) | 8 (88.9) | 8 (100.0) | 2 (100.0) |
| Plaque CT (HU) | 40±105 | 92±136 | 204±202 |
| ALT (U/L) | 21±12 | 34±30 | 30/21 |
| AST (U/L) | 22±8 | 30±25 | 25/26 |
| TAG (mM) | 1.7±1.2 | 1.4±0.67 | 2.4/- |
| TC (mM) | 3.9±0.85 | 3.4±0.43 | 5.2/- |
| HDL-C (mM) | 1.2±0.52 | 1.1±0.30 | 1.0/- |
| LDL-C (mM) | 2.0±0.75 | 1.7±0.28 | 3.3/- |

Gender is shown as number of patients and percentage, and all other values are mean ± SD. TAG, triacylglycerol; TC, total cholesterol; HDL, high-density lipoprotein cholesterol; LDL, low-density lipoprotein cholesterol.

Then the plaques were minced, homogenized, and subjected to density gradient centrifugation to isolate LDs, followed by differential centrifugation to separate LDs by size. The results verified that plaques contained few large LDs, with the majority being small LDs. The yellow coloration of the small LD fraction may reflect lipid oxidation within plaques (Figure 1D). Purified LDs showed a uniform spherical morphology under both fluorescence confocal microscopy and TEM (Figure 1E and 1F). The size of LDs was measured by Delsa Nano C Particle Analyzer, which showed that the average diameter of large LDs was about 1,600 nm and the average diameter of small LDs was about 300 nm (Figure 1G). The *in vitro* measurements were consistent with the *in situ* TEM observations of plaque LDs.

### Plaque lipid droplets are “fibrotic lipid droplets”

To characterize the proteins associated with LDs in atherosclerosis, we first analyzed various subcellular fractions of plaque homogenates. Silver staining revealed that the protein profiles were remarkably similar across all fractions (LD, cytosol, total membrane), a pattern distinct from that of typical tissues or cells. All fractions were dominated by two protein bands at approximately 40 kDa and 70 kDa (Figure 2A). We used Western blotting to validate these initial observations and assess the purity of our LD fraction. As expected, LD-resident proteins ADRP/PLIN2 and PLIN3 were mainly detected in the LD fraction. ACSL3 was also only detected in the LD fraction. Conversely, markers for potential contaminants like mitochondria (VDAC), the ER (Calnexin), and cytoplasm (GAPDH) were minimal, confirming the high purity of our isolation (Figure 2B). Critically, for the key lipases responsible for breaking down neutral lipids, NCEH1 (neutral cholesterol ester hydrolase) was rarely detected in the LD fraction (Figure 2B), and ATGL (adipose triglyceride lipase) was not detected in any plaque component, including the LD fraction (data not shown). This demonstrates that these LDs are metabolically inert.

**Figure 2.**
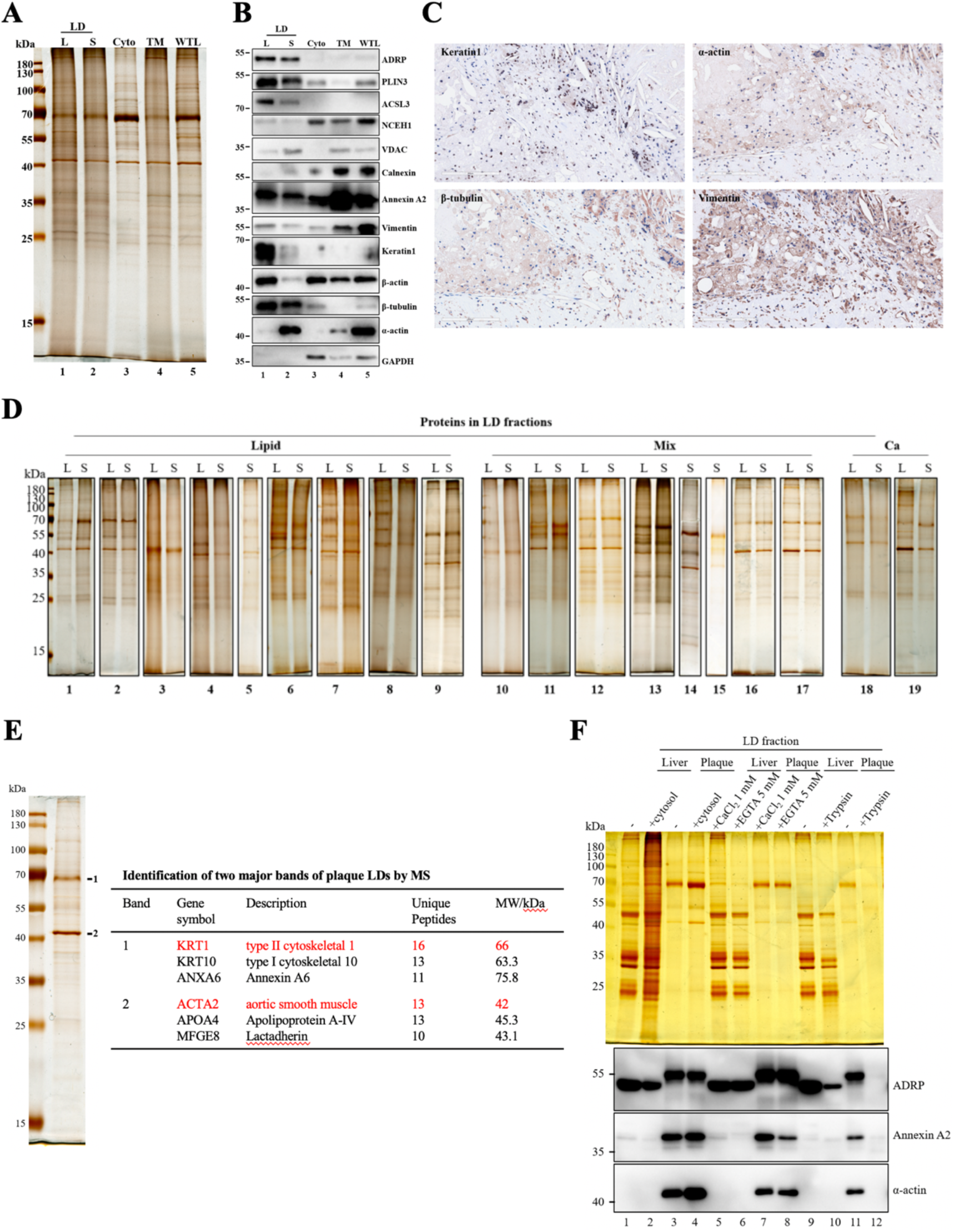
Protein profiles and *in vitro* characteristics of lipid droplets from human carotid artery plaques. A and B. Analysis and validation of proteins on plaque LD by silver staining and Western blot. Equal amounts of protein extracted from LD, cytosol (Cyto), total membrane (TM), and whole tissue lysate (WTL) were separated by SDS-PAGE, and the gel was used for silver staining as described in Materials and Methods or transferred to a membrane and were probed with indicated antibodies (include different cellular organelle markers and cytoskeletal proteins). C. Immunohistochemical staining of plaque cytoskeletal proteins in plaque sections. D. Silver staining analysis of LD proteins from 19 human plaque samples. L: large LDs; S: small LDs. E. MS identification of the two major bands of plaque LD proteins. F. *In vitro* recruitment assay of plaque LDs.

Surprisingly, plaque LDs contained multiple types of cytoskeletal proteins, including microfilament proteins α-actin and β-actin, tubulin protein β-tubulin, and intermediate filament proteins Vimentin and Keratin1 (Figure 2B). Immunohistochemical analysis further demonstrated that plaques contained abundant cytoskeletal proteins, which were predominantly enriched in foam-like cells within the plaques (Figure 2C). This crucial finding indicates that this dense cytoskeletal coat may be formed inside the foam cells before the LDs are released into the plaque core.

Having established the general fibrotic character, we focused on the purified LDs from the carotid artery plaques from all 19 patients. Silver staining of the LD proteins revealed a remarkably consistent profile across all plaques of the patients, and between the large and small LD fractions from each patient, which was dominated by two major bands at approximately 40 kDa and 70 kDa (Figure 2D). This high degree of similarity differs from the protein profiles of LDs purified from other tissues or cells, or from those observed under different pathological conditions.

To identify the prominent proteins in plaque LDs, the two major protein bands observed in the silver-stained LD protein gel were excised and subjected to proteomic analysis. The two bands were identified as containing abundant cytoskeletal proteins, Keratin 10/1 and ACTA2, respectively (Figure 2E), consistent with the Western blotting results (Figure 2B).

We next performed *in vitro* experiments to probe the properties of these unique LDs. Purified plaque LDs were incubated with plaque cytosol to see if they could interact with soluble proteins. At the same time, LDs from normal mouse liver were used as controls, with liver LDs incubated with liver cytosol. The results showed that LDs of normal liver could recruit multiple cytosolic proteins but could not recruit actin. In contrast, plaque LDs were able to recruit additional α-actin from the cytosol (Figure 2F, lanes 1-4), indicating the presence of specific actin-binding proteins or lipids on the surface of plaque LDs. Annexins are Ca^2+^-regulated membrane binding and actin-interacting proteins, potentially mediating actin binding to LDs^[23]^. As shown in Figure 2B, abundant annexin A2 was detected in plaque LDs and cytosol. Therefore, we investigated the role of the calcium-dependent annexin pathway in affecting actin binding to plaque LDs. The results revealed that annexin A2 relied on calcium ions for LD binding while neither the addition of calcium nor a calcium chelator (EGTA) affected actin binding to plaque LDs (Figure 2F, lanes 5-8). This suggests the plaque LD-actin interaction is not mediated by annexins. Furthermore, we treated the LDs with trypsin to determine whether the LD surface cytoskeleton was exposed or encapsulated within a membrane. The results demonstrated that when the plaque LDs were treated with trypsin, the α-actin was readily digested, similar to the LD protein PLIN2 /ADRP (Figure 2F, lanes 9-12). This demonstrates that the actin shell is not encapsulated within a membrane but is externally exposed. These findings reveal that the fibrotic coat is a dynamic surface, capable of further interacting with cytoskeletal elements in the plaque environment.

### Proteomic profiles of lipid droplets and tissue lysates from atherosclerotic plaques

To gain deeper insight, we performed a comprehensive, quantitative proteomic analysis on LDs and their corresponding whole tissue lysates (WTL) from plaques classified as lipid-rich (LIP), mixed (MIX), and calcified (Ca) (Table 2). For LD proteins, the LIP group included 7 samples (D_AS1–7), the MIX group included 3 samples (D_AS8–10), and the Ca group included 2 samples (D_AS11–12). For whole tissue lysate (WTL) proteins, the LIP group contained 7 samples (W_AS1–7), the MIX group included 4 samples (W_AS8–11), and the Ca group contained 1 sample (W_AS12). Using data-independent acquisition (DIA) mass spectrometry, we constructed a deep proteome of these LDs and plaques. After normalization of the proteomic data, shared proteins of LD fractions from the LIP, MIX, and Ca groups contained 4078, 2599, and 2926 proteins, respectively (Figure S1A). In comparison, the WTL fractions contained 4643, 4644, and 5678 proteins, respectively (Figure S1B).

**Table 2.**
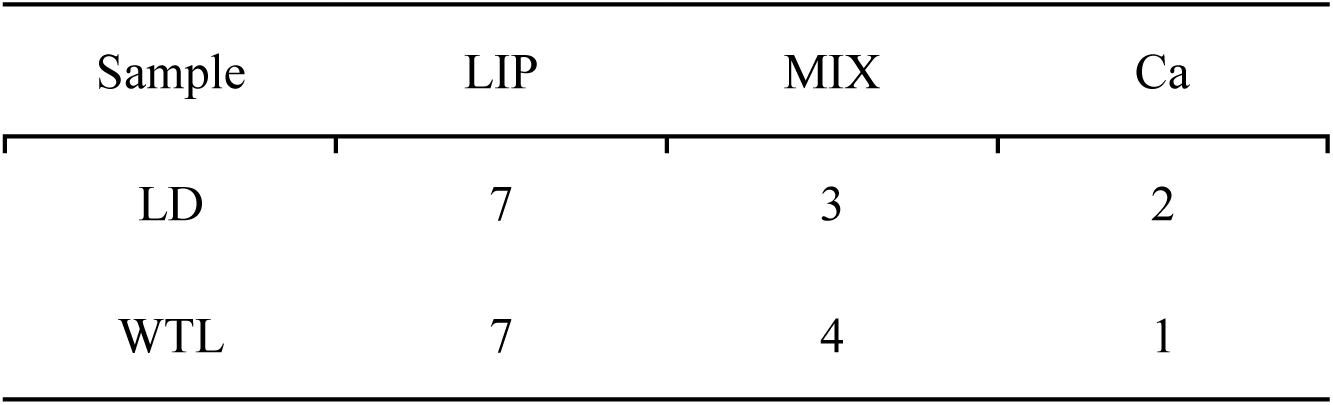
Sample Grouping for LC-MS/MS Analysis.

| Sample | LIP | MIX | Ca |
| --- | --- | --- | --- |
| LD | 7 | 3 | 2 |
| WTL | 7 | 4 | 1 |

The LD fractions shared 2312 proteins across all groups, while the WTL fractions shared 4254 proteins (Figure 3A). These shared proteins across all groups were further sorted by subcellular localization and known functions based on literature and NCBI resources. In the plaque LD proteome, secretory proteins are the main proteins, accounting for about 48.7% of the total protein, followed by cytoskeletal proteins and cytoskeletal regulatory proteins, accounting for about 19.3% and 5.1% respectively. This proteomic overview reinforces the identity of plaque LDs as “fibrotic” organelles, structurally dominated by cytoskeletal elements. In addition, LD proteins account for 1.8% (Figure 3B, panel a). The major LD proteins identified by MS are shown in Table 3. We compared them with LD proteins from other tissues or cells and listed the proteins unique to plaque LDs or THP-1 foam cell LDs in Table 4. In the WTL proteome, the main categories were secretory proteins, cytoskeletal proteins, cytoskeletal regulatory proteins and cytosol proteins, accounting for 55.4%, 2.7%, 7.6% and 4.7% respectively. LD proteins account for 0.1% (Figure 3B, panel b), confirming the success and necessity of our LD purification for studying these organelles. This suggests that plaques are primarily composed of abundant secretory proteins. KEGG pathway enrichment analysis was also performed on shared proteins identified in all LD and WTL samples. The most enriched pathway in both datasets including “Complement and coagulation cascades”, “Endocytosis”, “phagosome”, “Cytoskeleton in muscle cells” and “Regulation of actin cytoskeleton”, supporting the fibrotic identity of plaque LDs. Notably, “Cholesterol metabolism” was significantly enriched in the LD proteome compared to WTL, suggesting a tighter connection between LD and cholesterol metabolism (Figure 3C).

**Figure 3.**
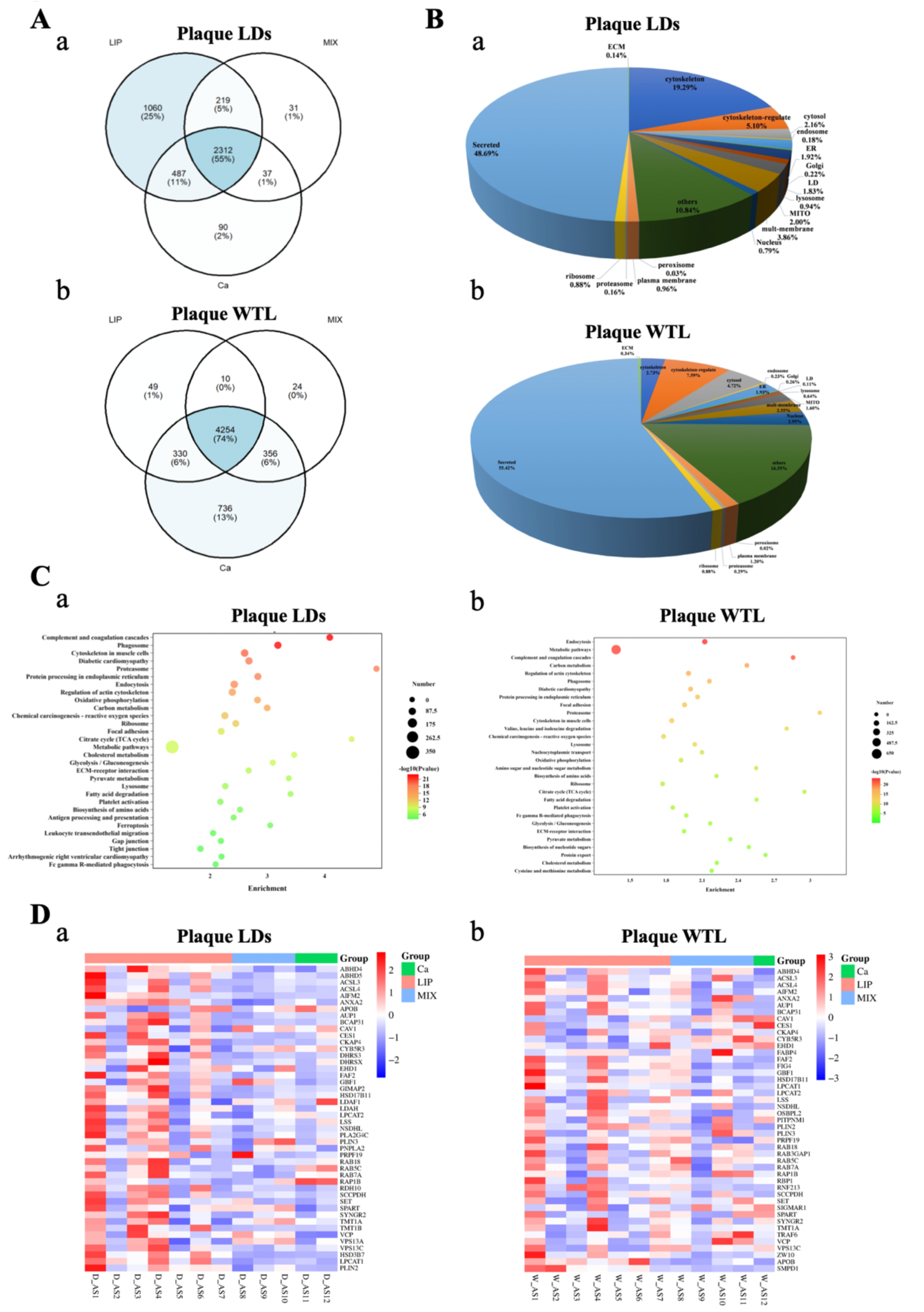
Proteomic analysis of plaque lipid droplets and tissue lysate. A. Venn diagram of the shared proteins in three types of plaques. a. Venn diagram of proteins commonly identified in three types of plaques in LD samples. b. Venn diagram of proteins commonly identified in three types of plaques in WTL samples. B. Subcellular localization of the shared plaque LD and WTL proteins. Based on literature and NCBI online resources, the shared LD and WTL proteins in all plaque samples identified by LC-MS were classified according to subcellular distribution and known functions. a. Subcellular localization of the shared plaque LD proteins. b. Subcellular localization of shared plaque WTL proteins. C. KEGG pathway enrichment analysis of shared proteins identified in LD and WTL samples. D. Heatmap of LD-resident proteins in three types of plaque. Heatmap showing the abundance of LD-resident proteins identified in LD and WTL samples, screened based on GO cellular component (GO_CC) enrichment analysis. LIP, lipid-rich plaque, MIX, mixed plaque; Ca, calcified plaque.

**Table 3.**
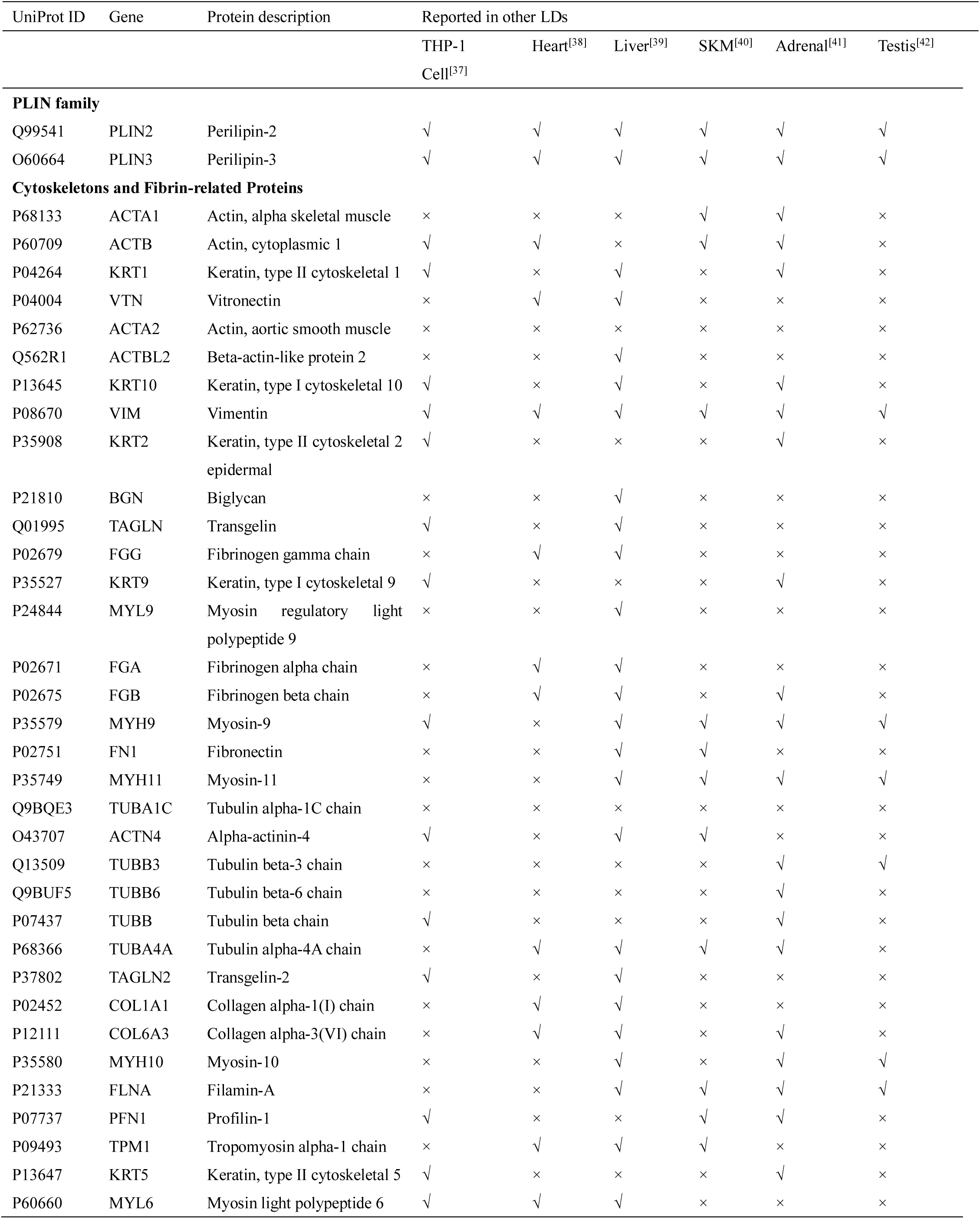
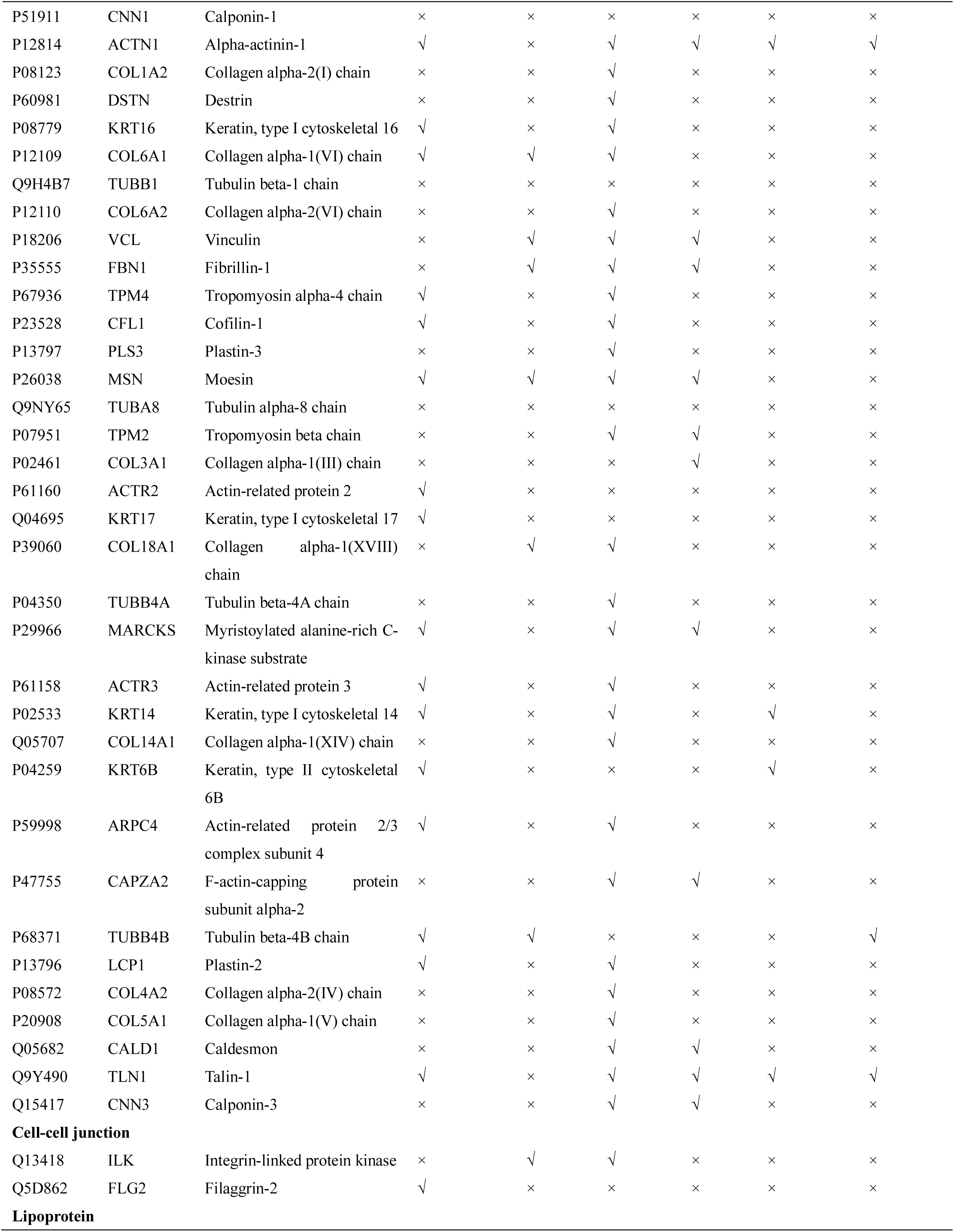
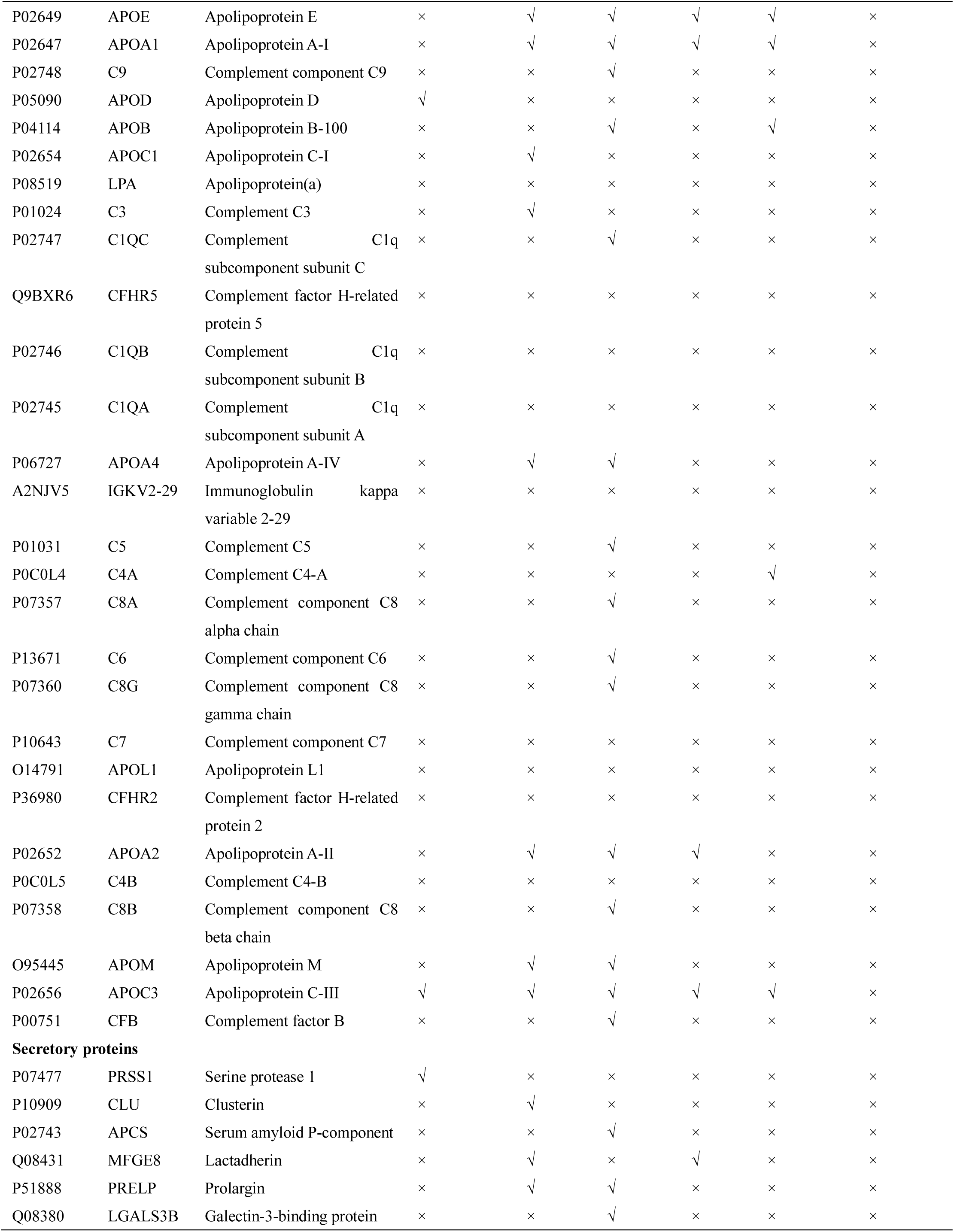
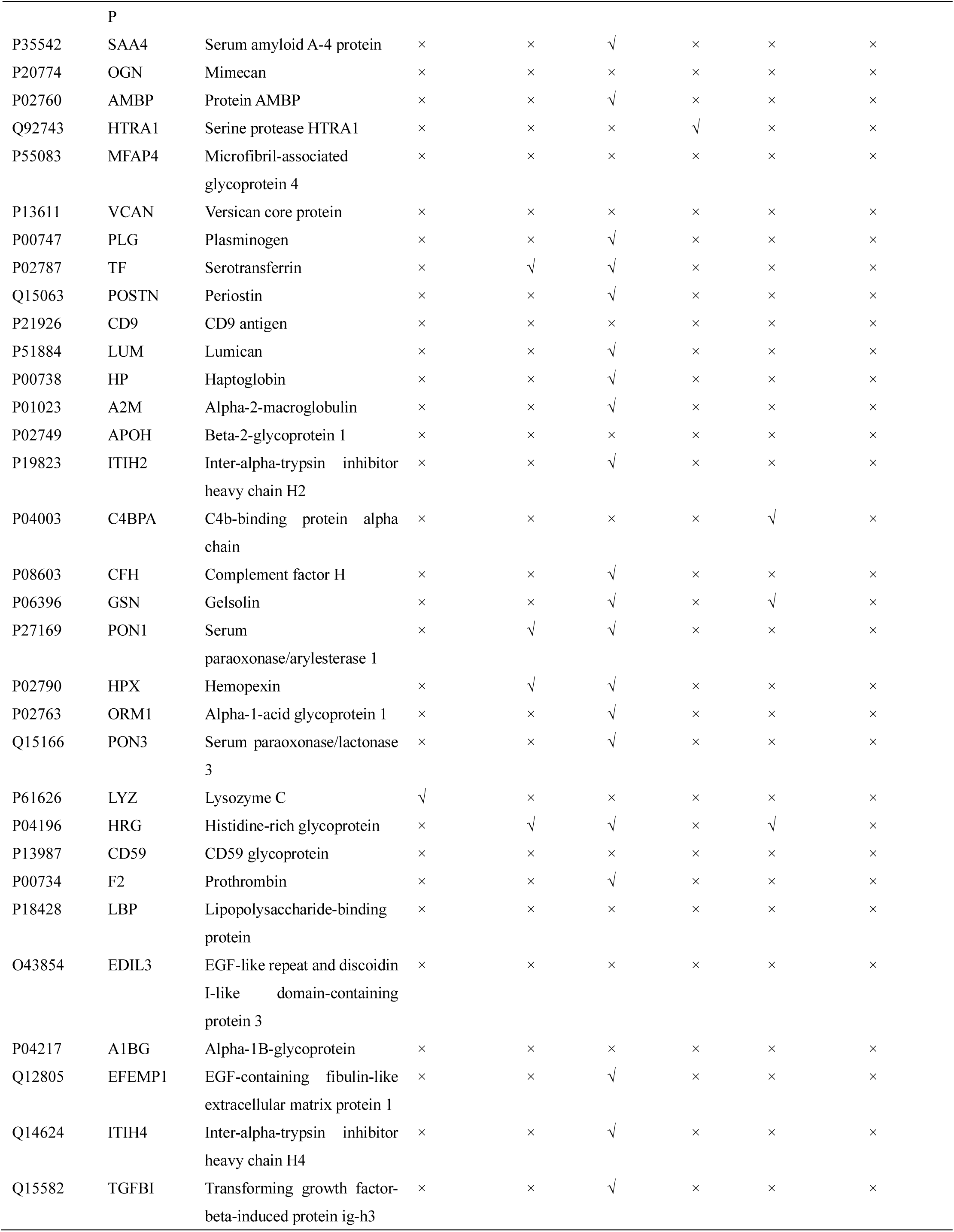
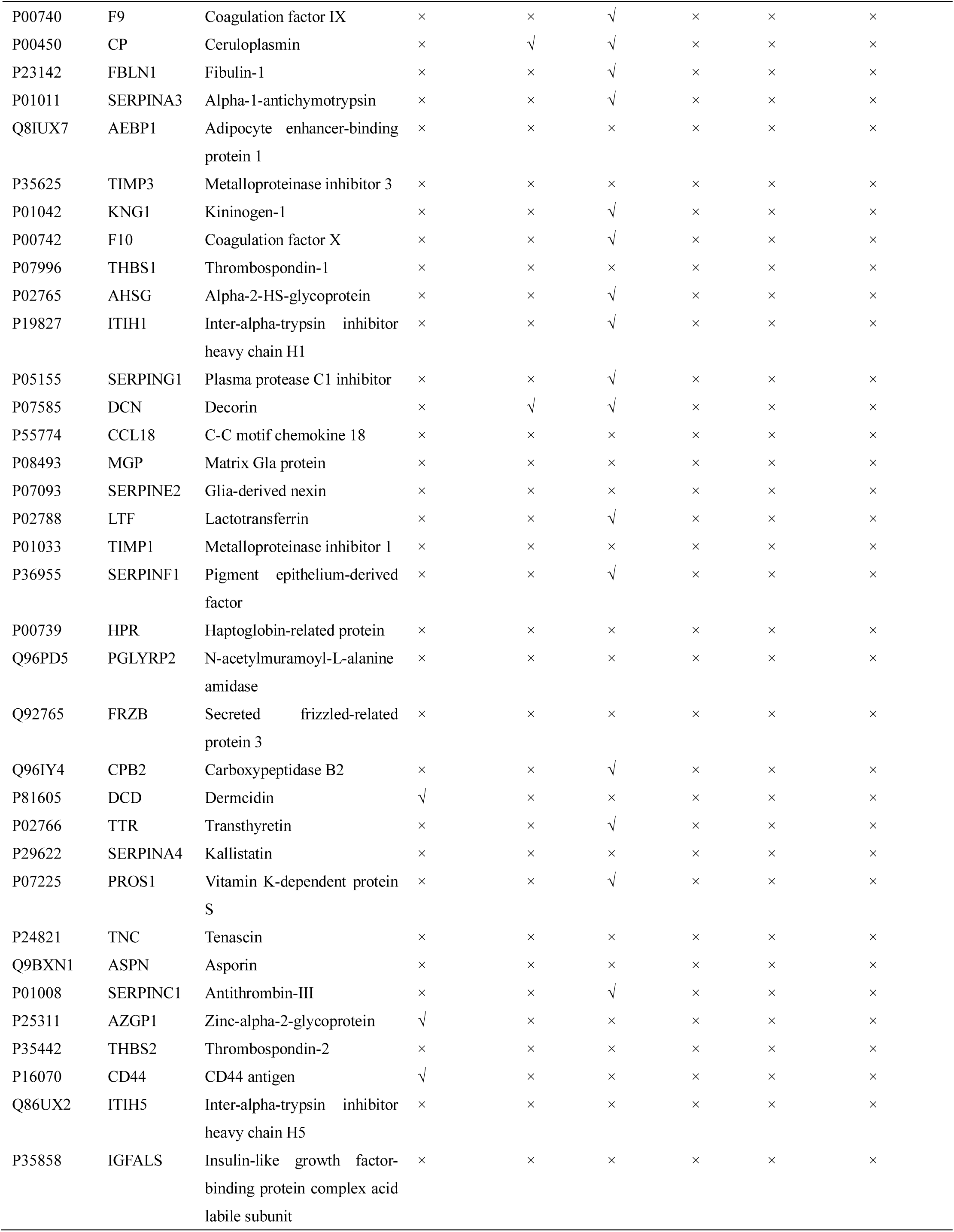
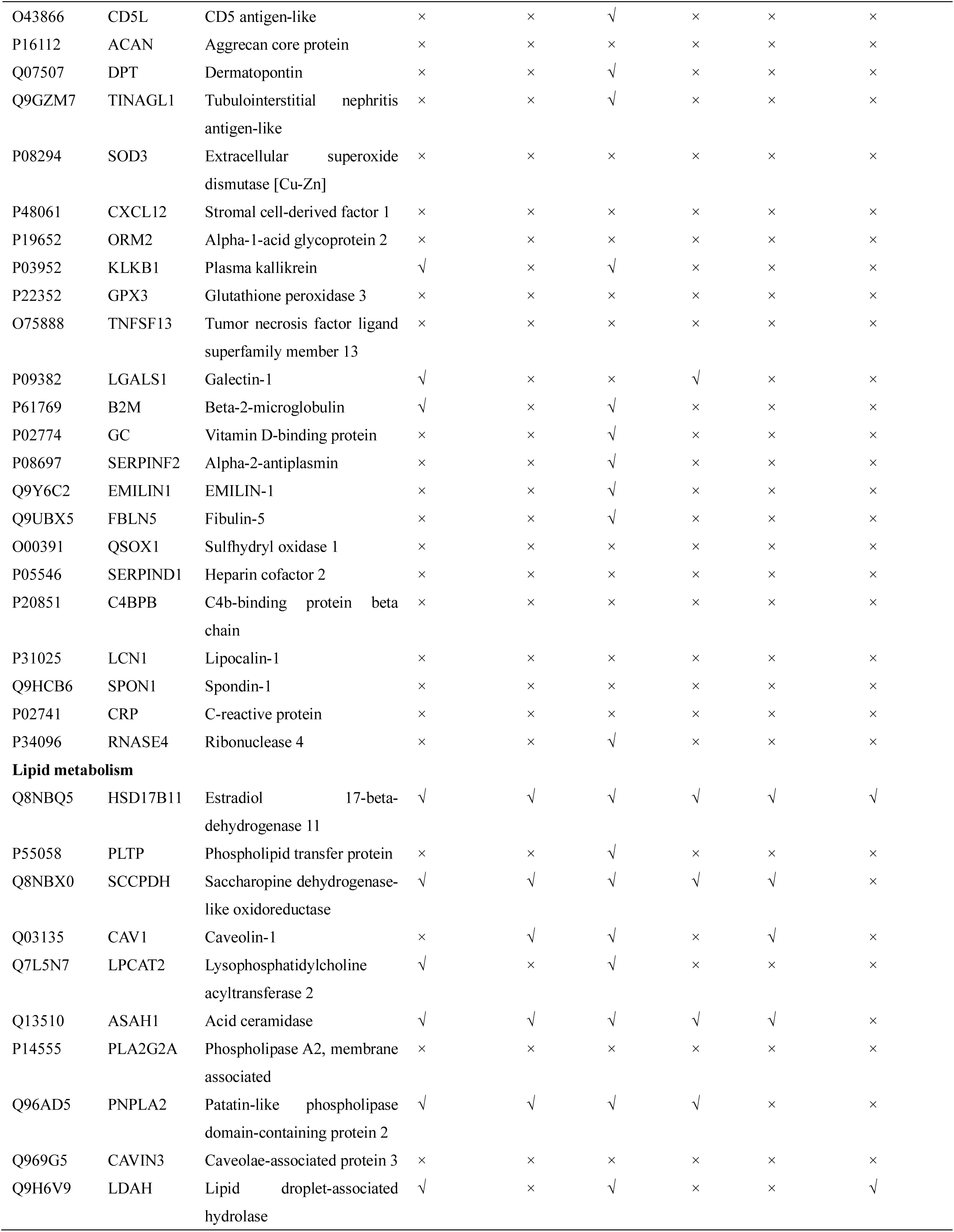
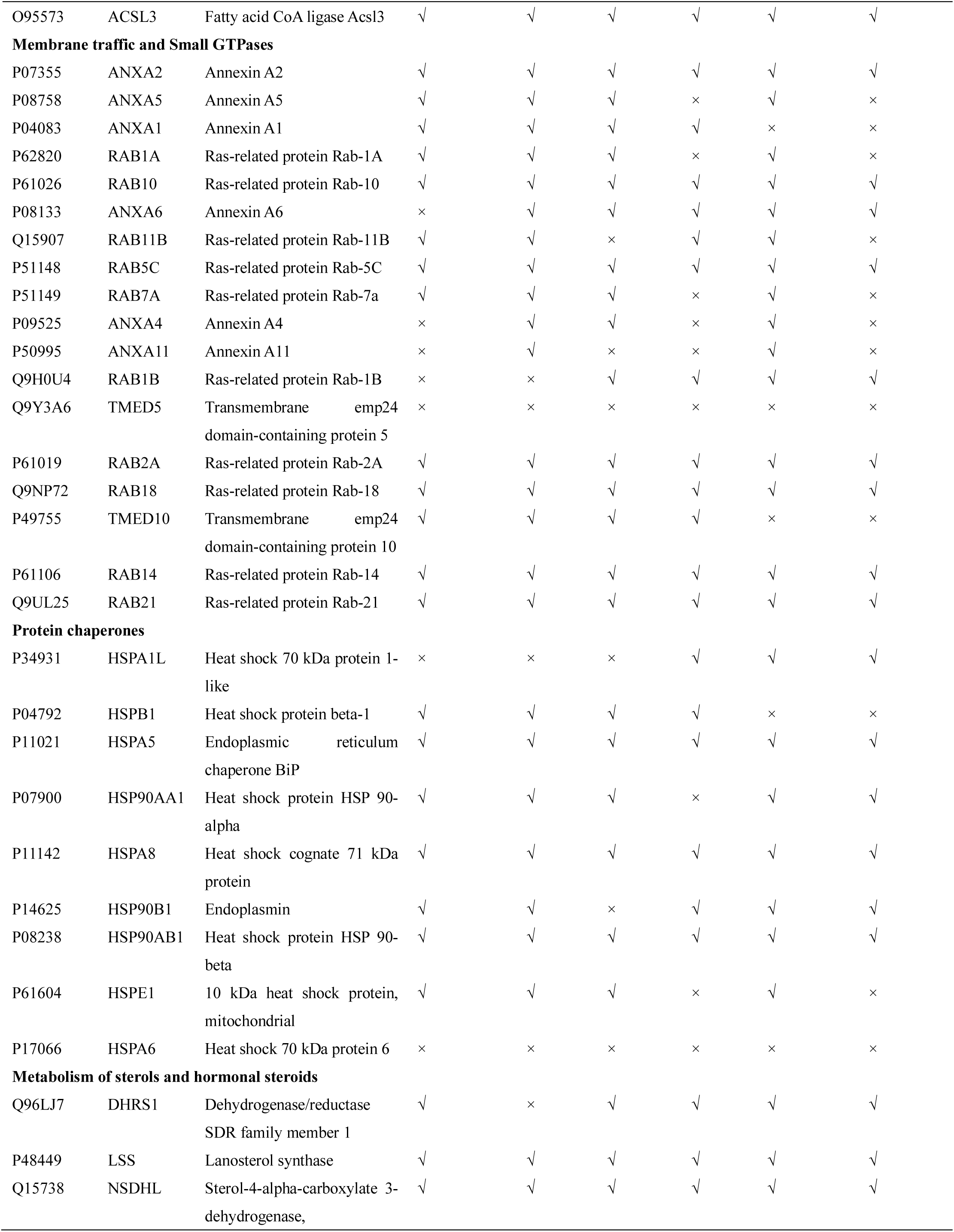
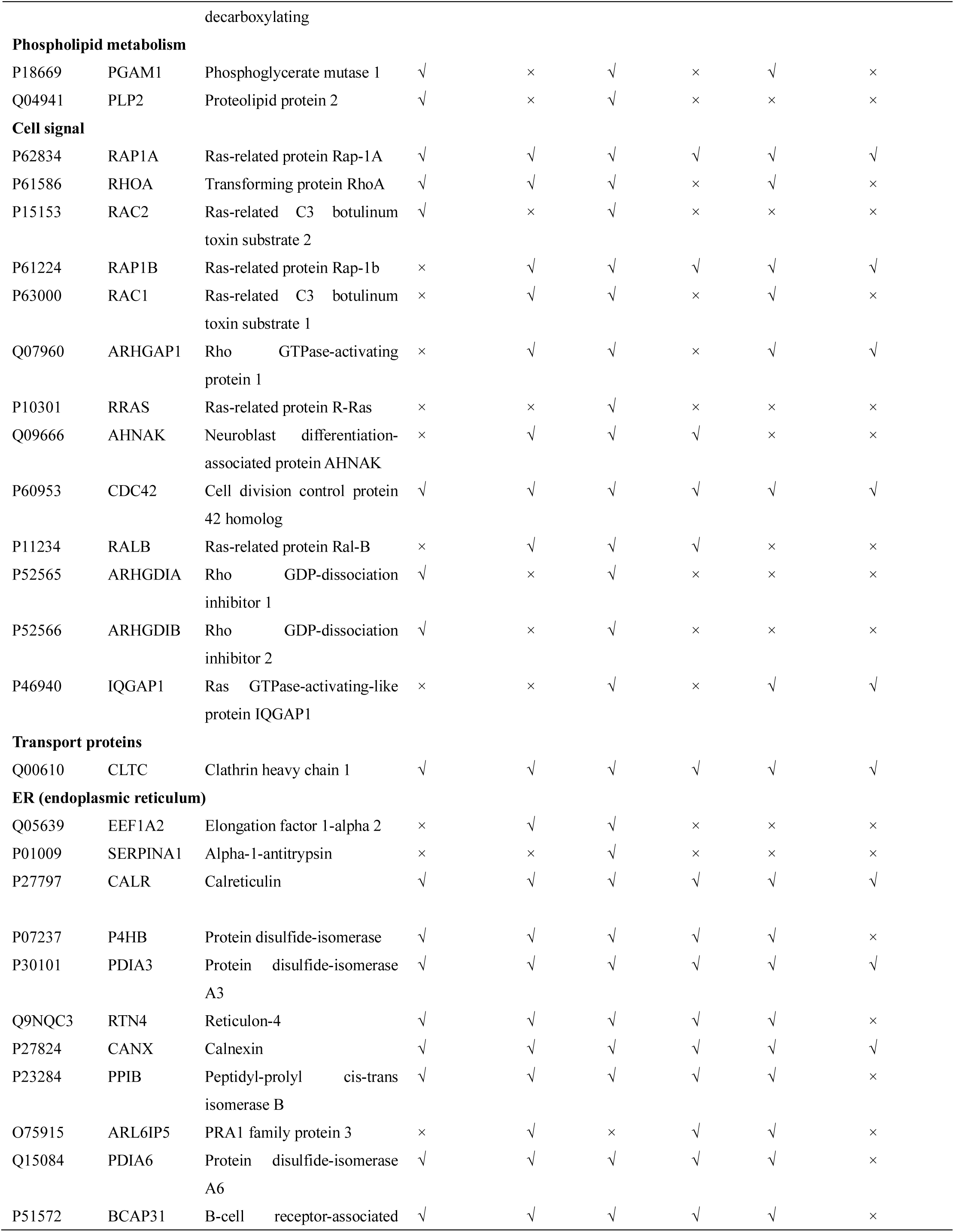
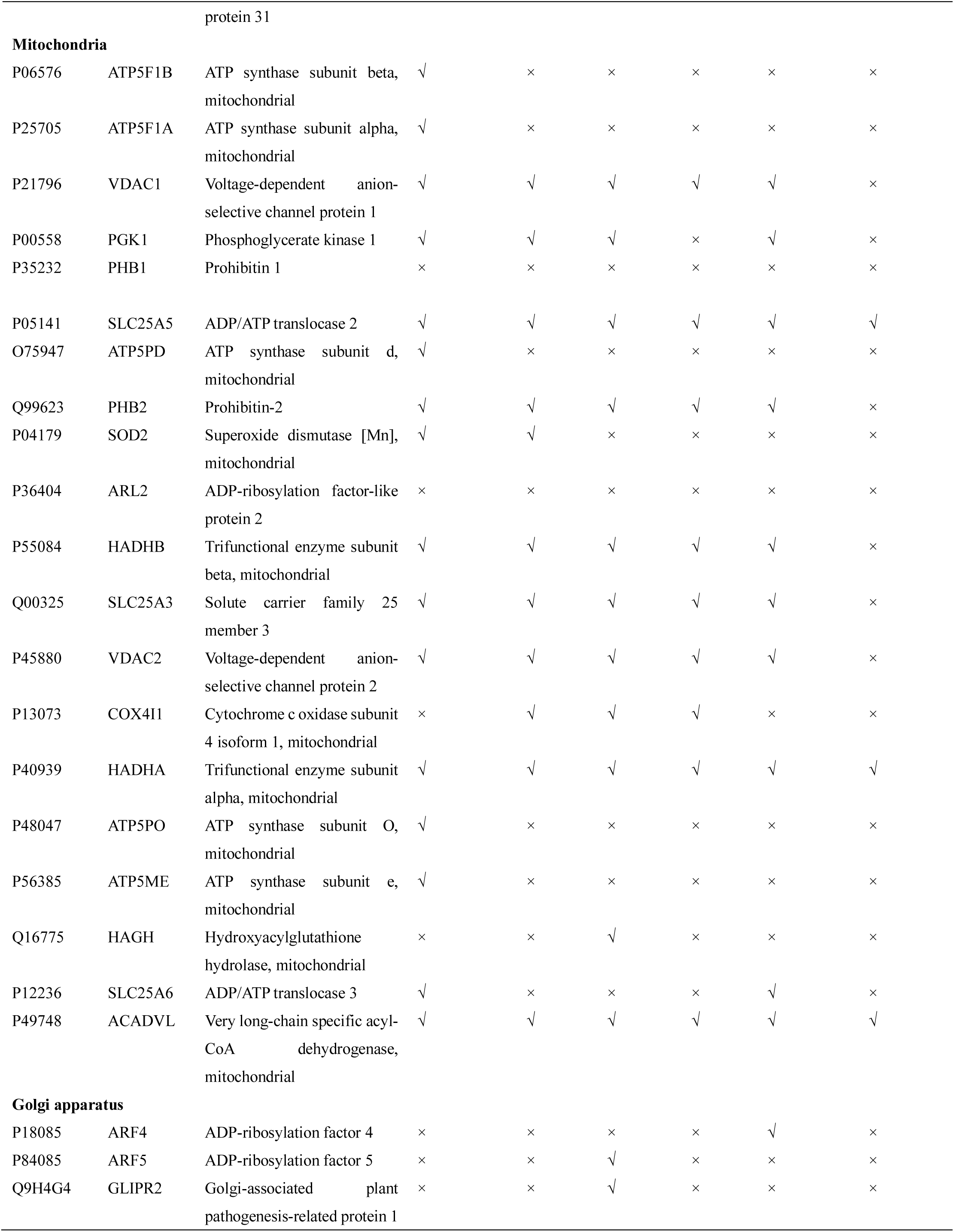
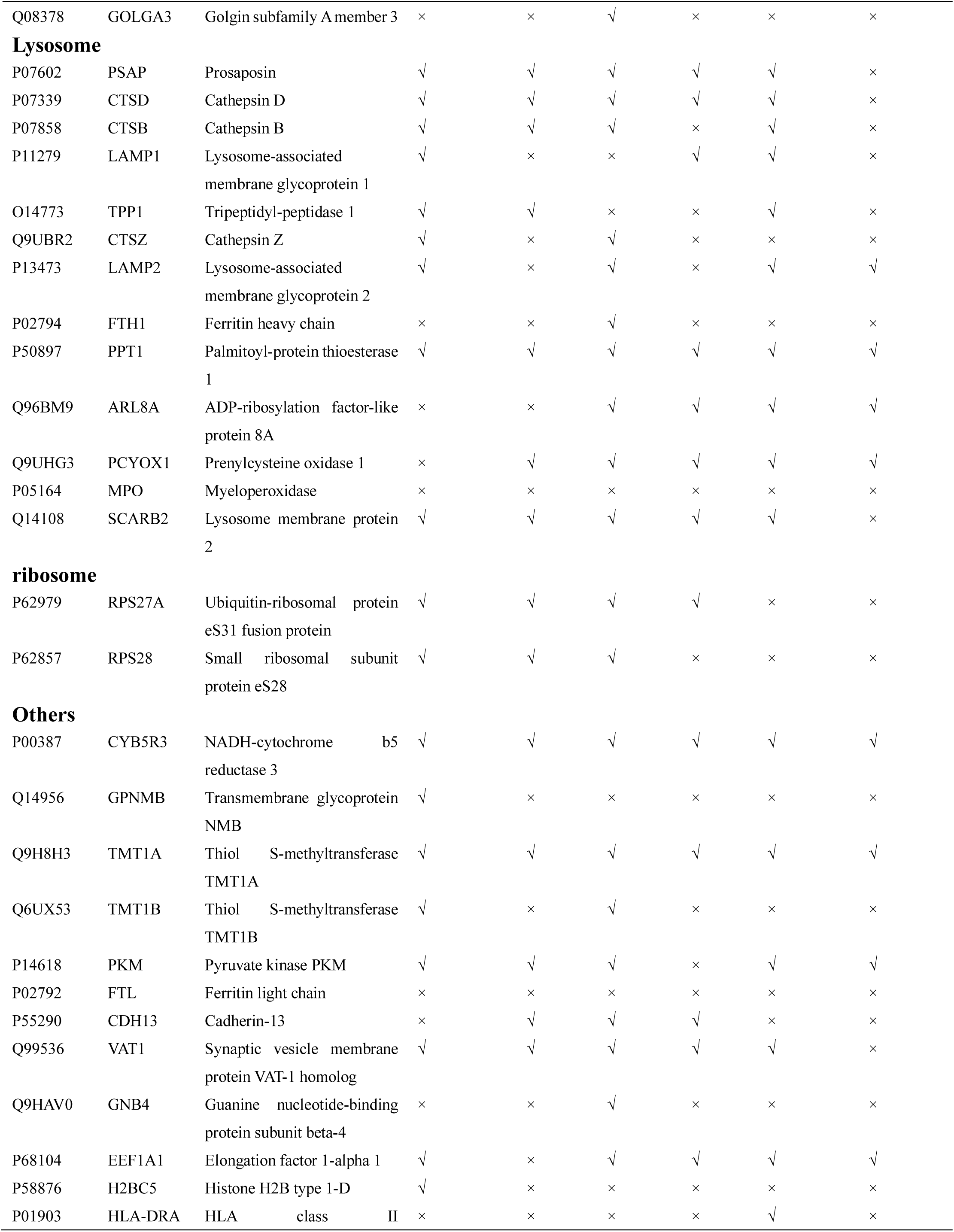
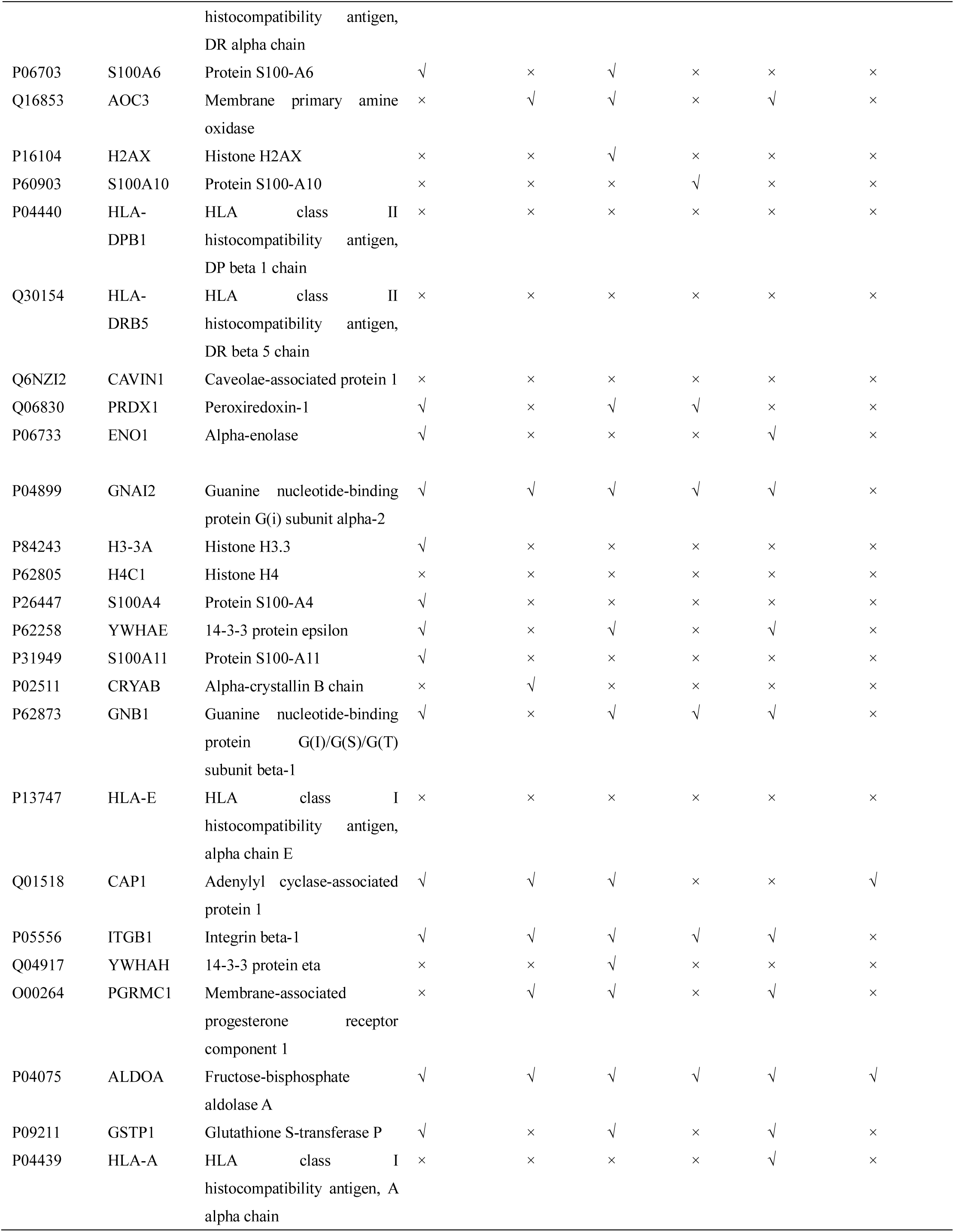
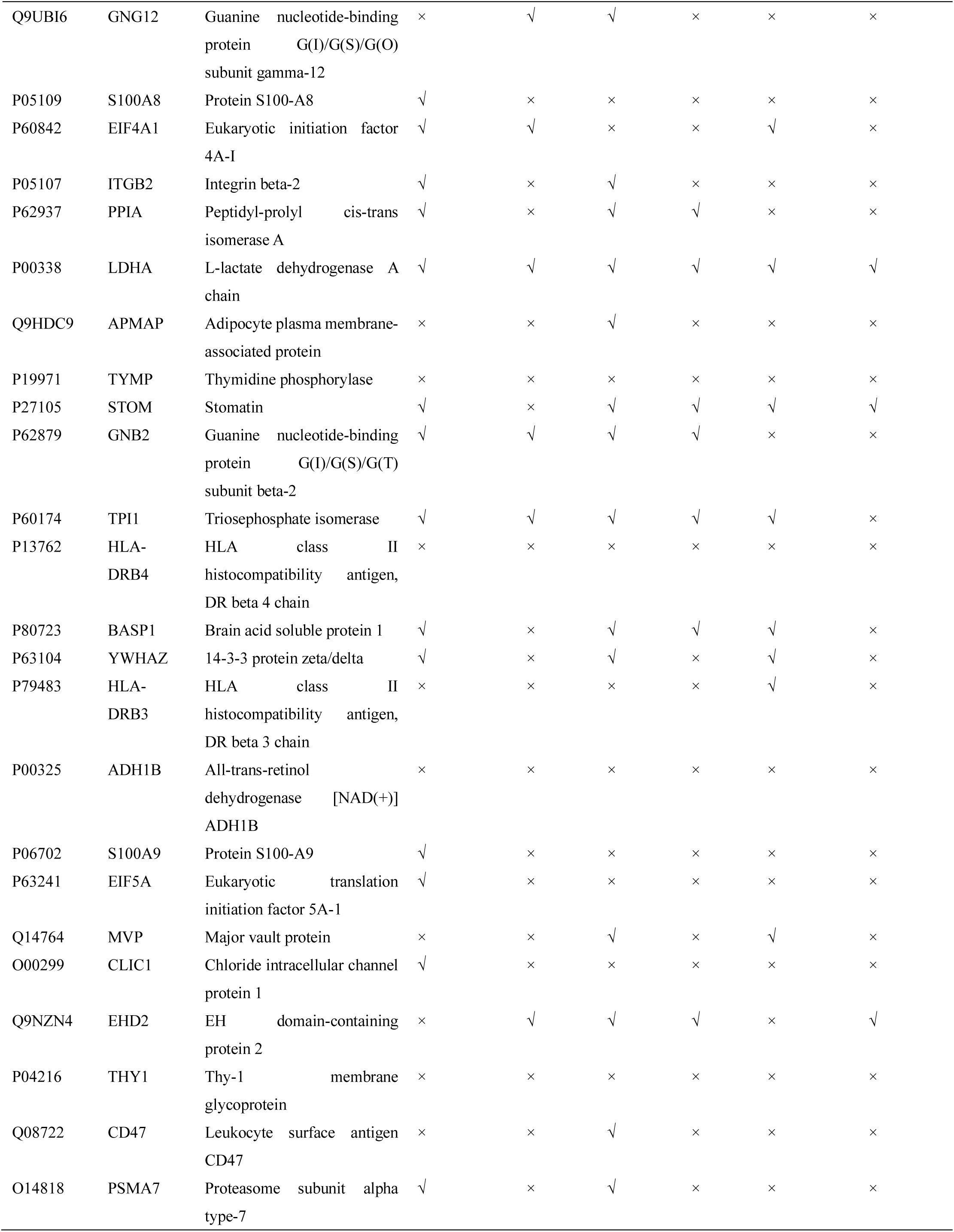
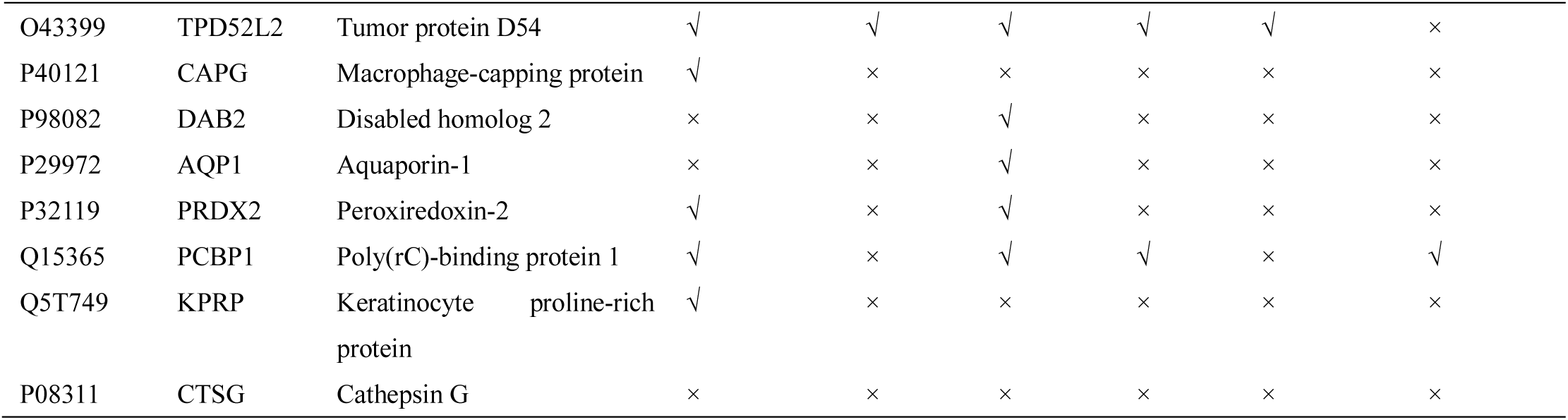
The main plaque LD proteins identified by MS.

**Table 4.** Specific proteins identified in plaques or THP-1 foam cell LDs.

| Specific protein in plaque LDs | Specific protein in plaque and THP-1 foam cell LDs |
| --- | --- |
| ACTA2 TUBA1C CNN1 TUBB1 TUBA8<br>APOL1 MFAP4 VCAN APOH CD59 LBP<br>EDIL3 A1BG AEBP1 TIMP3 THBS1<br>CCL18 MGP SERPINE2 TIMP1 HPR<br>PGLYRP2 FRZB SERPINA4 TNC ASPN<br>THBS2 ITIH5 IGFALS ACAN SOD3<br>CXCL12 ORM2 GPX3 TNFSF13 QSOX1<br>SERPIND1 C4BPB LCN1 SPON1 CRP<br>PLA2G2A CAVIN3 TMED5 HSPA6<br>ATP5F1B ATP5F1A ATP5PO ATP5ME<br>MPO OGN FTL HLA-DPB1 HLA-DRB5<br>CAVIN1 H4C1 HLA-E TYMP PHB1<br>HLA-DRB4 ADH1B THY1 CTSG | ACTR2 KRT17 FLG2 APOD LYZ DCD<br>AZGP1 PRSS1 GPNMB H2BC5 H3-3A<br>S100A4 S100A11 S100A8 S100A9 EIF5A<br>CLIC1 CAPG KPRP |

We next investigated how the proteome changes with plaque progression from early-stage lipid-rich (LIP) plaques to late-stage, non-LIP (MIX/Ca) plaques. Principal component analysis (PCA) of the LD and WTL datasets revealed a clear separation between the LIP group and the non-LIP groups in the LD samples, whereas the WTL samples showed no obvious separation among the three groups (Figure 4A). This suggests that LD-associated proteins correlate better with plaque classification compared to WTL proteins. This demonstrates that the protein composition of the LDs is a more sensitive and specific indicator of the plaque’s pathological state than the global tissue proteome. Further supporting this, GO analysis showed that LIP plaques, often called “soft plaques,” had a significantly higher content of LD-resident proteins (Figure 3D), consistent with a more active state of lipid handling in these earlier-stage lesions. Analysis of proteins annotated as cholesterol metabolism-related showed no significant differences between the groups (Figure S1C).

**Figure 4.**
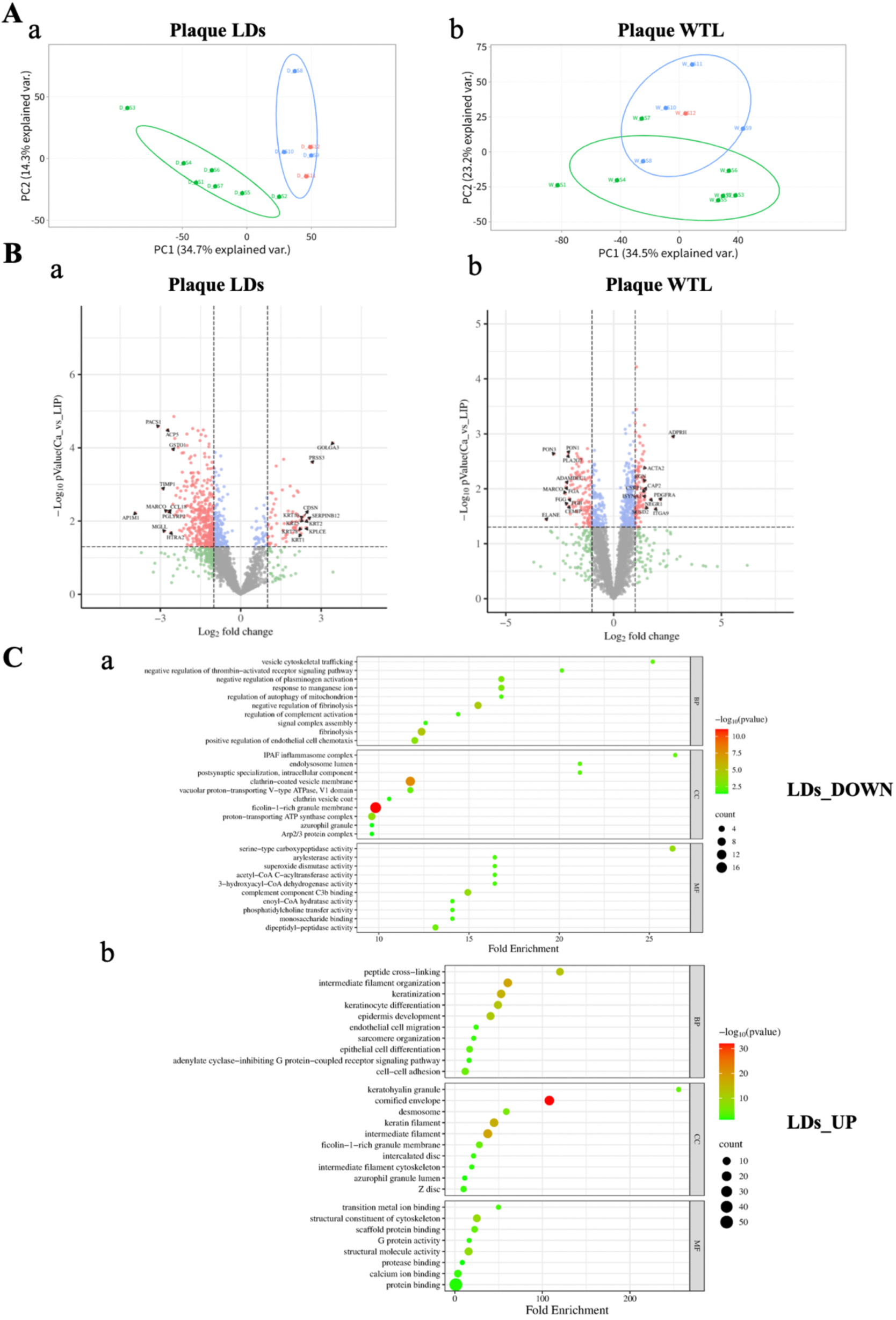
Differential analysis of the proteomes of distinct plaque types. A. Principal component analysis (PCA) of LD and WTL samples. a. PCA plot of all 12 LD samples, b. PCA plot of all 12 WTL samples. B. Top 10 differentially expressed LD and WTL proteins between LIP and non-LIP groups. C. GO enrichment analysis of differentially expressed LD proteins between LIP and non-LIP groups.

To identify the specific proteins driving the separation observed in the PCA, we performed a differential expression analysis between the LIP and non-LIP groups. We combined the data of MIX and Ca groups into a single non-LIP group based on PCA results. In the LD samples, comparison between the LIP and non-LIP groups identified 444 differentially expressed proteins (DEPs), of which 62 proteins were upregulated and 382 were downregulated in the non-LIP group relative to the LIP group. In the WTL samples, a total of 205 DEPs were detected between the LIP and non-LIP groups, including 90 upregulated and 115 downregulated proteins in the non-LIP group relative to the LIP group. These results are visualized in volcano plots (Figure 4B). In the LD samples, 197 proteins were exclusively detected in the seven LIP samples but absent in the five non-LIP samples. In contrast, in the WTL samples, only one protein was uniquely detected in all seven LIP samples but not in the five non-LIP samples. Notably, no proteins were identified that were present in five non-LIP samples but absent in seven LIP samples for either LD or WTL. These unique proteins were then combined with the downregulated DEPs in non-LIP group for subsequent enrichment analyses. GO enrichment analysis revealed that the downregulated proteins in LD samples of non-LIP group were predominantly metabolic enzymes (Figure 4C, panel a), whereas in WTL samples they were mainly proteins associated with lipoprotein binding or lipid transport activity (Figure S2A, panel a). KEGG pathway enrichment analysis further showed that downregulated proteins in both LD and WTL samples of non-LIP group were significantly enriched in cholesterol metabolism-related pathways (Figure S2B and S2C, panel a). Conversely, upregulated proteins in both LD and WTL samples were mainly cytoskeletal proteins (Figure 4C and S2A, panel b). The upregulated proteins in the LD samples were relatively few and primarily associated with cytoskeletal regulation, which is consistent with the GO results (Figure S2B, panel b). In the WTL samples, upregulated proteins were mainly involved in amino acid metabolism pathways (Figure S2C, panel b).

Taken together, these findings strongly suggest a functional transformation of LDs during atherosclerosis. As plaques advance and calcify, the LDs transition from being metabolically active organelles to becoming inert, fibrotic structures through the systematic replacement of their metabolic enzymes with a dense structural shell.

### Plaque lipid droplet core is dominated by an unprecedented abundance of polyunsaturated cholesteryl esters

Further, we analyzed the neutral lipid core that this fibrotic LDs encases. TLC analysis showed that the plaque LDs were mainly composed of CEs, with an average content of 75%±9%, while TAGs were a minor component at 13%±8% (Figure 5A and 5B). While the exact ratio varied slightly between patients, CE dominance was a universal feature. This composition, though high in CE, is notably different from the ∼95% CE reported by LANG *et al*. in 1970^[17]^.

**Figure 5.**
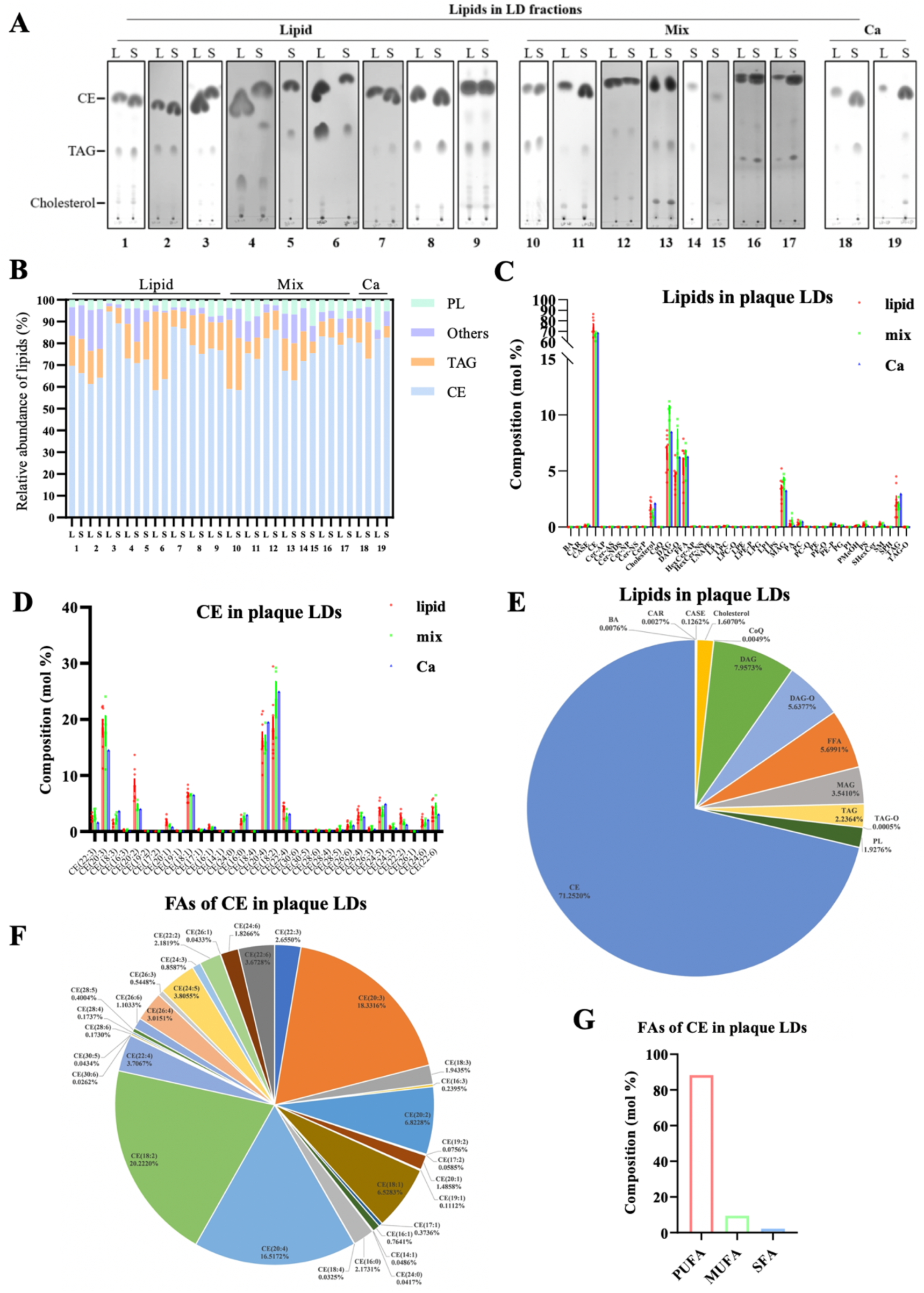
Lipidomic analysis of carotid artery plaque lipid droplets. A. Thin layer chromatography (TLC) analysis of lipid composition of plaque LDs from 19 plaque samples. B. Quantitative analysis of TLC lipid bands. Image J quantitative analysis of the proportion of LD cholesterol ester (CE), triacylglycerol (TAG), phospholipids (PL) and other lipids from 19 plaque LDs in Figure 5A. C. Mole percentage of lipids in LDs of three types of plaques through lipidomic analysis. D. Overall lipid composition of plaque LDs. The mole percentages of different lipid represent the average of a total of 13 LD sample. E. Mole percentages of CE species in LDs of three types of plaques. F. Overall CE species composition of plaque LDs. The mole percentages of different CE species represent the average of a total of 13 LD sample. G. The ratio of CE containing different fatty acid types.

Then to define the lipid composition of the plaque LDs, we performed a detailed lipidomic analysis on 13 representative samples of purified small LDs. A total of 42 lipid species, including CE, diacylglycerides (DAG), ether-linked DAG (DAG-O), free fatty acids (FFA), monoacylglycerides (MAG), and phospholipids (PL) (including phosphatidylcholine (PC), PA, sphingomyelin (SM) and PE-P classes), were characterized. The composition of different lipid species is similar in lipid, mixed, and calcified plaques (Figure 5C). Consistent with the TLC results, the plaque LD core was dominated by CE, accounting for 71.3%±7.2% (molar ratio) of total lipids. As the overall lipid composition was relatively consistent across the 13 plaque LD samples, we summarized the average molar percentage of each lipid species in these samples (Figure 5D). In addition to CE, DAG accounted for 8.0%±2.2%, DAG-O accounted for 5.6%±1.9%, FFA accounted for 5.7%±1.7%, monoacylglycerides (MAG) accounted for 3.5%±1.0%, TAG accounted for 2.2%±1.0%, and cholesterol accounted for 1.6%±0.6% (Figure 5C). TAG, DAG and DAG-O within the LDs were composed primarily of common saturated, monounsaturated fatty acids and diunsaturated fatty acid (e.g., 16:0, 18:0, 18:1, 18:2) (Figure S3, A-D). The free fatty acids (FFA) in LDs are also mainly 16:0 and 18:0 (Figure S3E).

Surprisingly, the fatty acid composition of these CEs was striking. A total of 33 CE species were identified. Most CE containing PUFA (including 20:3, 18:2, 20:4, 22:3, 18:3, 20:2, 22:4, 26:4, 24:5, 24:6, and 22:6) (Figure 5, E and F). The three most abundant species were those containing LA (18:2), DGLA (20:3), and the key pro-inflammatory precursor arachidonic acid (AA, 20:4). The three CE species accounted for 20.2%±5.8%, 18.3%±4.1%, and 16.5%±3.3% of the total CE, respectively (Figure 5E). In contrast, CEs containing the most common fatty acids in plasma, palmitic (16:0) and oleic (18:1) acid, were only minor components, accounting for 2.2%±0.6% and 6.5%±0.9% of the CE content, respectively. Among the 71.3% CE, 63% were PUFA-CE, 8.3% were CE with saturated and monounsaturated side chains. Therefore, 88.3% of all CE species were polyunsaturated (PUFA-CEs), 9.5% were monounsaturated fatty acid (MUFA-CE) and 2.2% were saturated fatty acid (SFA-CE) (Figure 5G), a level of enrichment not typically seen in other tissues. Crucially, this extreme PUFA enrichment was exclusive to CE.

For plaque LD phospholipids, PC accounts for 18.0%±7.8% of the phospholipids in plaque LDs, PA accounts for 16.8%±16.2%, PE-P accounts for 13.1%±3.0%, SM accounts for 11.9%±6.2%, PS accounts for 11.1%±7.1%, and the rest includes a small amount of phosphatidyl methanol (PMeOH), phosphatidyl glycerol (PG), lysophosphatidylcholine (LPC) and other different phospholipids (Figure 6, A and B). The molar percentages of PA, PE-P, SM, and PS in plaque LD phospholipids all exceeded 10% (Figure 6B). The high abundance of sphingomyelin, a precursor to the pro-atherogenic signaling molecule ceramide, suggests these LDs could be a platform for harmful signaling. Furthermore, phosphatidylethanolamine (PE) accounts for only 0.3%±0.1% of the total phospholipids in LD, while the PE-P is striking enrichment. PE-P is a class of plasmalogens reported to be involved in inflammation and antioxidant activities^[24,25]^, which seems to be related to the high content of PUFA including AA in LDs (Figure 5, E and F). The fatty acid side chains in PL are mainly 16:0, 18:1 and 18:0, accounting for 32.4%, 13.8% and 9.0% respectively (Figure 6C). Phospholipids are mainly composed of saturated fatty acid side chains, with SFA, PUFA, and MUFA accounting for 53.1%, 23.6%, and 23.3%, respectively (Figure 6D). In addition, PC was mainly PC (16:0_18:2) and PC (16:0_18:1), SM was mainly SM (d18:1_16:0), PA was mainly PA (16:0_22:5), and PE-P was mainly PE (P-15:0_20:3) (Figure S4, A-D). Together, instead of being dominated primarily by PC, the plaque LD phospholipid displayed a complex and functionally significant composition.

**Figure 6.**
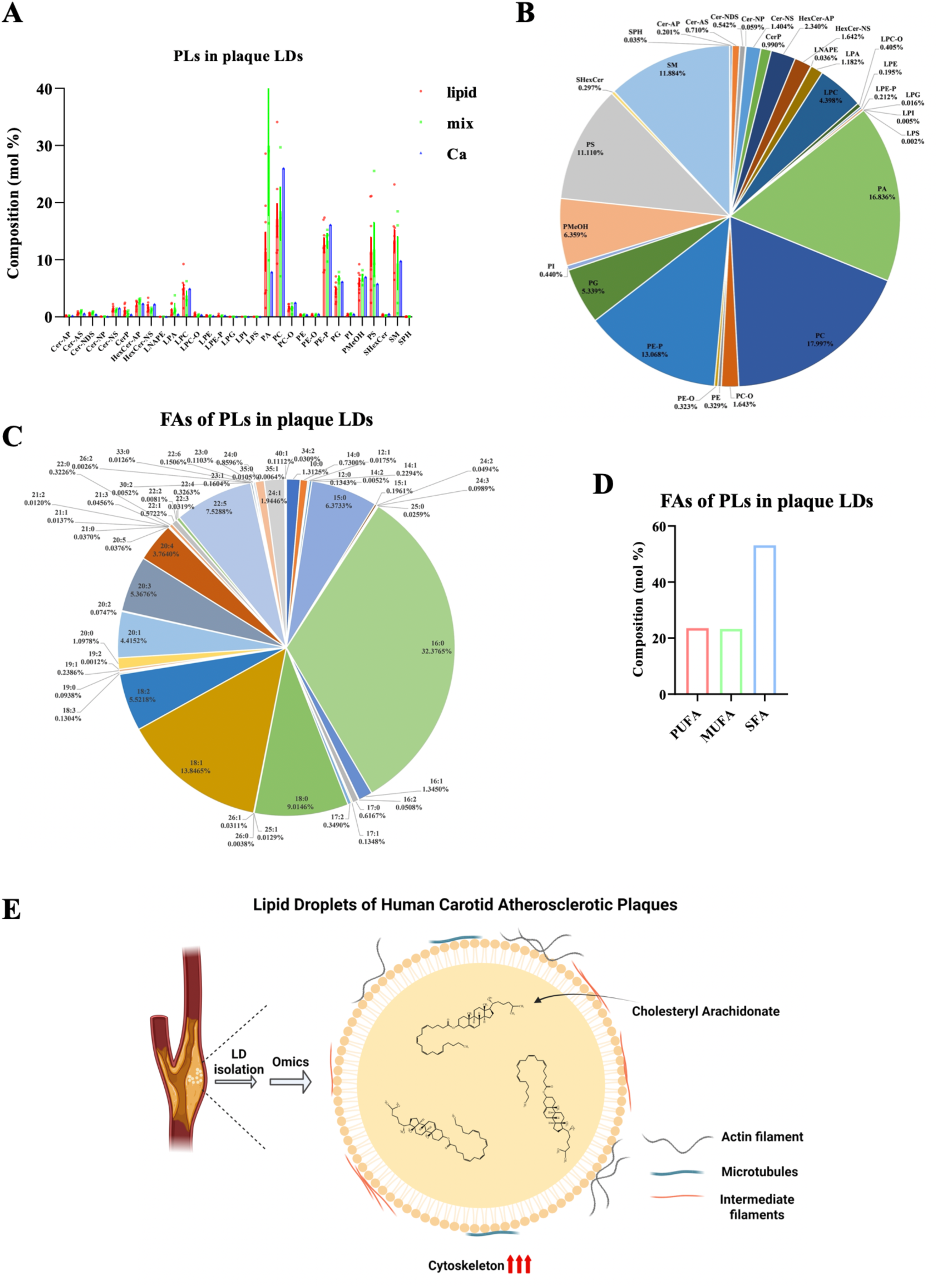
Analysis of phospholipid composition in lipid droplets from carotid artery plaques. A. Mole percentages of phospholipid (PL) species in LDs of three types of plaques. B. Overall PL species composition of plaque LDs. The mole percentages of the different PL species represent the average of a total of 13 LD sample. C. Overall fatty acid composition of PL in plaque LDs. The mole percentages of different fatty acid composition of PL represent the average of a total of 13 LD sample. D. The ratio of PL containing different fatty acid types. E. Schematic diagram of LD characteristics in human carotid atherosclerotic plaques.

## Discussion

Atherosclerosis is driven by the accumulation of LDs in plaques, yet the specific nature of these organelles is not fully understood. Investigating the progression of atherosclerosis at the LD level provides critical insights for understanding this pathological process. Our results showed that the LDs in human plaques were mainly small LDs with a diameter less than 500 nm, and there are few LDs larger than 1 µm in the plaque (Figure 1C). This may be related to the fact that the plaque LDs are rich in CE. It has been reported that CE-rich LDs are easy to crystallize, while TAG-rich LDs will not crystallize and can form µm-sized LDs^[26]^. It has also been reported that the presence of other lipids such as TAG in LDs facilitates the clearance of CE. Therefore, the accumulation of abundant CE also makes this type of LDs difficult to clearance^[27]^. In this study, we successfully isolated LDs from different types of human atherosclerotic plaques and performed the first in-depth quantitative proteomic and lipidomic analyses of these organelles. A key discovery of our study is that plaque LDs are encased in a dense coat dominated by cytoskeletal proteins, with mass spectrometry identifying Keratin 10/1 and ACTA2 as its main components (Figure 2, D and E). Moreover, proteomic results show that the content of cytoskeletal proteins in plaque LD proteins accounts for 19.3% (Figure 3B, panel a). And TEM observation of plaque LDs also found that there are many fibrous structures next to LDs (Figure 1C). All the data show that plaque LDs are encased by fibrotic structure. This fibrotic structure likely provides the stability for these LDs to persist as extracellular bodies in plaques. Interestingly, while the overall composition of large and small LDs is similar (Figure 2D), subtle differences in their cytoskeletal protein profiles (Figure 2B) suggest they may originate from distinct sources or function differently within the plaque.

Crucially, our *in vitro* experiments revealed that this coat is not a static, encapsulated barrier. It is an externally exposed and dynamic surface capable of recruiting additional α-actin from the cytosol. The most critical consequence of this coat is the creation of a metabolically inert organelle. Its dense structure physically excludes key lipases like NCEH1, effectively trapping cholesterol esters inside by preventing their hydrolysis, despite the presence of these enzymes elsewhere in the plaque (Figure 2B).

Notably, fibrotic LDs that recruit more actin suggest that cytoskeletal proteins on the surface of plaque LDs may be gradually formed by the enhanced interaction with LDs in the cells, especially during the formation of foam cells. This is supported by our observation of abundant cytoskeleton in foam cells of plaques (Figure 2C). Similarly, the enhanced interaction between the cytoskeleton and LDs in the formation of CE-rich LDs has also been reported^[28,29]^ However, the precise mechanism and functional significance of these interactions remain unclear, even though our data indicate that they occur independently of the calcium–annexin pathway.

The core of these fibrotic LDs is as unusual as their surface proteins. Our lipidomic analysis revealed an unprecedented enrichment of polyunsaturated fatty acid-cholesteryl esters (PUFA-CEs), which accounted for 88.3% of all CEs. This enrichment is highly specific, as other neutral lipids like TAGs and the free fatty acid pool were composed of standard saturated and monounsaturated fats (Figure 5F and S3B), which suggests that foam cells in human plaques may prefer to synthesize PUFA-CE. This may be related to the unique spatial structure of cholesterol. Cholesterol is a planar structure, while LDs are spherical. Therefore, when CE accumulates in LDs and lacks neutral lipids such as TAG to fill other gaps in LDs, the side chains of cholesterol need to be flexible fatty acids rich in double bonds rather than saturated fatty acids to fill the gaps. Of course, this requires further experimental proof.

Two of the most significant component of this PUFA pool is AA (20:4) and its direct precursor, DGLA (C20:3) (Figure 5F). AA is closely related to inflammatory response, and its metabolites such as PGE2 (prostaglandins E2) can promote acute inflammatory response. In active macrophages, lipid hydrolases remain on the surface of LDs during the lipid accumulation process. These hydrolases release the stored AA, which is metabolized by catalytic enzymes oxygenases: cyclooxygenase (COX), lipoxygenase (LOX), and cytochrome P450 into a spectrum of bioactive mediators including PGE2^[30]^. These bioactive mediators may further attract macrophages, causing the plaque to gradually expand. This finding provides a direct molecular link between plaque LDs and the chronic inflammation that drives atherosclerosis, and it helps explain the therapeutic efficacy of COX inhibitors like aspirin, which can reduce inflammation and alleviate the development of atherosclerosis^[31–33]^. Therefore, AA-enriched LDs in macrophages or foam cells may be core to the inflammatory response. Moreover, it will be critical in future studies to determine if AA is also enriched in large LDs of plaque. Furthermore, the high concentration of DGLA (20:3) may present a new area of research to investigate its metabolism and potential role in generating other bioactive lipid mediators within the plaque.

We also identified a substantial amount of SM in the phospholipids of plaque LDs (Figure 6, A and B). Ceramide, a bioactive metabolite of SM, has been reported to promote the development of atherosclerosis^[34–36]^. This suggests that LDs within atherosclerotic plaques may also metabolize SM to generate ceramide, thereby contributing to plaque formation.

The differential proteomic analysis provides a window into how these unique organelles evolve with the disease. Proteomic analysis of plaque LDs indicate that, compared to the LIP group, the non-LIP group exhibits a decrease in metabolic-related proteins and an increase in cytoskeletal proteins (Figure 4C and S2A-C). This suggests that the transformation of plaques from “soft plaques” to “hard plaques” after calcification may be a continuous process. This process may effectively reinforce the fibrotic coat over time, further locking the AA-rich lipid core in its metabolically inert state and preventing the resolution of its inflammatory potential. This dynamic change in LD composition appears to be a key feature of plaque maturation and hardening.

This study provides the first comprehensive omics analysis of carotid plaque LDs. However, several limitations should be acknowledged, and important future directions remain to be explored. The LD preparations likely contain a mixture of LDs from within intact foam cells and those from the extracellular necrotic core. Future work should aim to separate and independently analyze these LD populations. It is possible that the large LDs, which were too scarce in our samples for omics analysis, primarily originate from intact foam cells, while the more numerous small LDs constitute the extracellular pool. Additionally, due to the limited availability of patient samples, our analysis of “hard” calcified plaques was based on only two cases, requiring us to combine them with the “mixed” plaque group. We are currently collecting more samples from calcified plaques to strengthen this comparison. Future studies will help elucidate, in greater detail, how distinct LD subpopulations contribute to the progression of atherosclerosis.

In conclusion, our work defines the human atherosclerotic plaque LDs as a unique “bad” organelle, characterized by a dynamic, fibrotic coat that confers metabolic inertia and a lipid core that serves as a massive reservoir for inflammatory precursors (Figure 6E). In the future, the mechanisms that reveal how the fibrotic coat is assembled should be investigated. Besides, to explore the strategy how to disrupt the coat may be a viable way to reduce the inflammatory burden of plaque and the development of atherosclerosis.

## Supplemental Data

**Figure S1.**
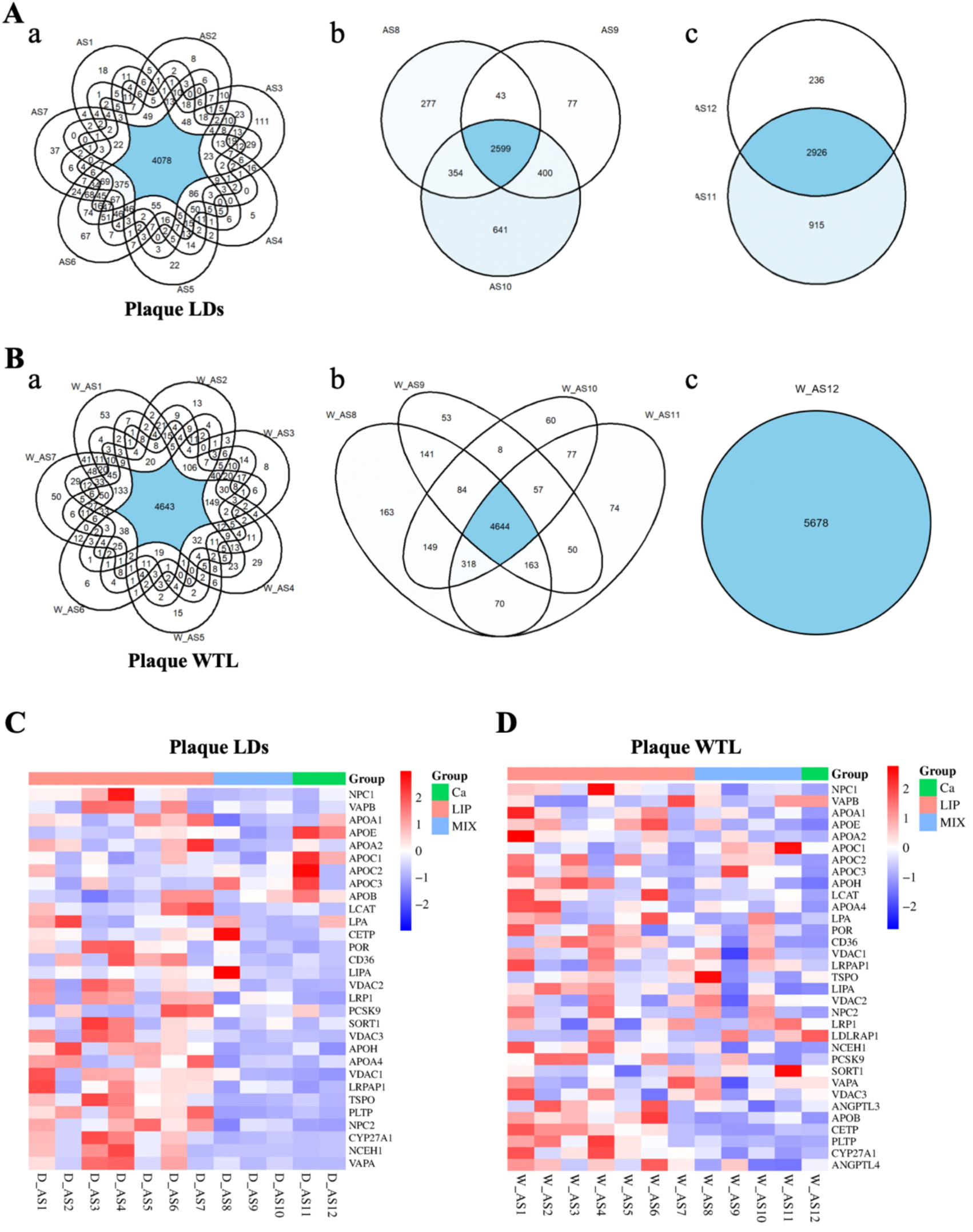
Venn diagrams and heatmaps for protein identification by MS. A. Venn diagrams of the overlap of proteins identified by LC-MS/MS in LD samples from different plaque types. a. Venn diagram of shared proteins identified in seven LIP samples, b. Venn diagram of shared proteins identified in three MIX samples, c. Venn diagram of shared proteins identified in two Ca samples. B. Venn diagrams of the overlap of proteins identified by LC-MS/MS in WTL samples from different plaque types. a. Venn diagram of proteins identified in all seven LIP samples, b. Venn diagram of proteins identified in all four MIX samples, c. Venn diagram of proteins identified in two Ca samples. C. Heatmap of cholesterol metabolism–related proteins in three types of plaque. Heatmap showing the abundance of cholesterol metabolism–related proteins identified in LD and WTL samples, screened based on KEGG pathway enrichment analysis.

**Figure S2.**
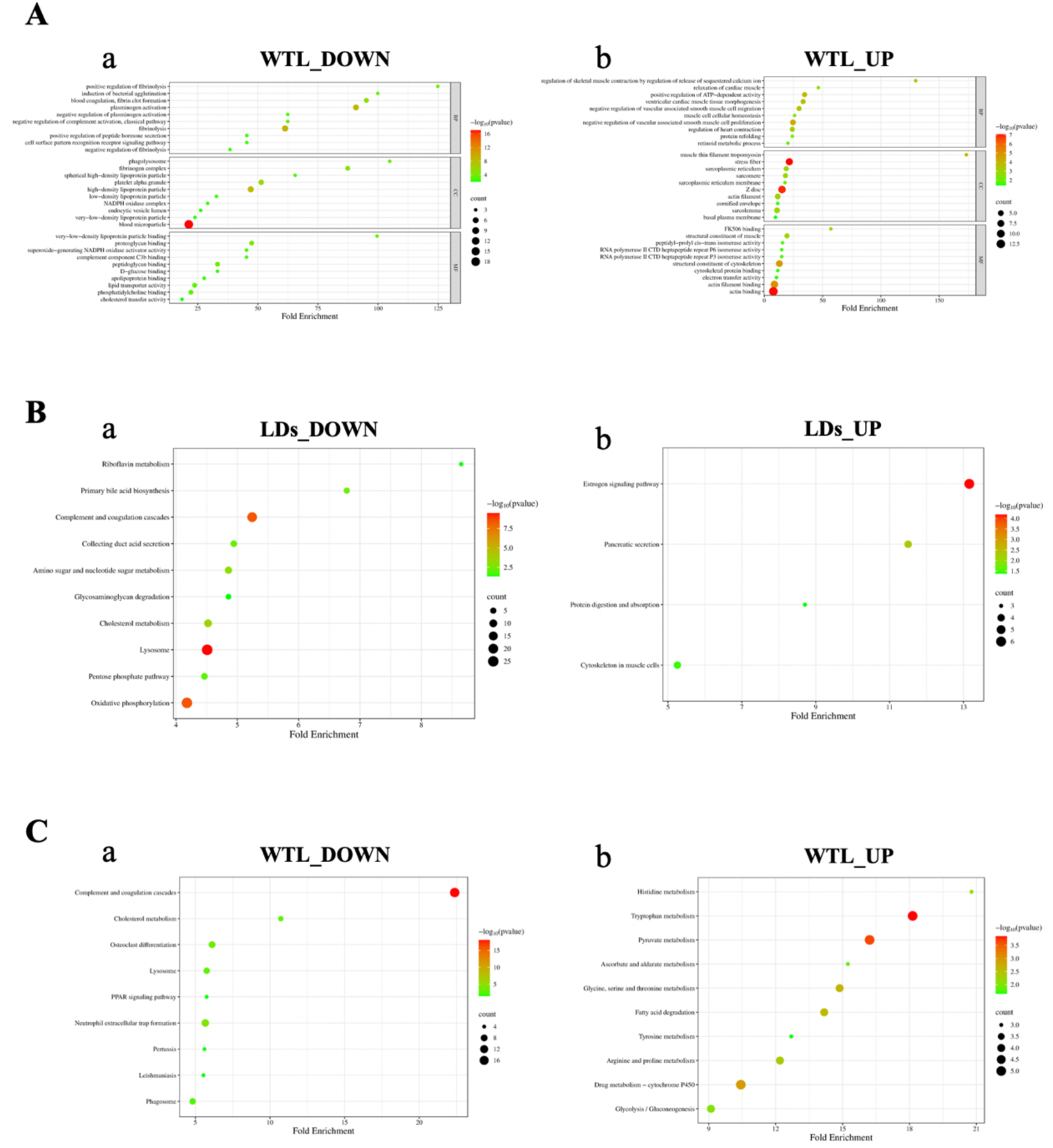
GO and KEGG analysis of the proteomes of different types of plaques. A. GO enrichment analysis of differentially expressed proteins in WTL samples. B and C. KEGG pathway enrichment analysis of differentially expressed proteins in LD and WTL samples.

**Figure S3.**
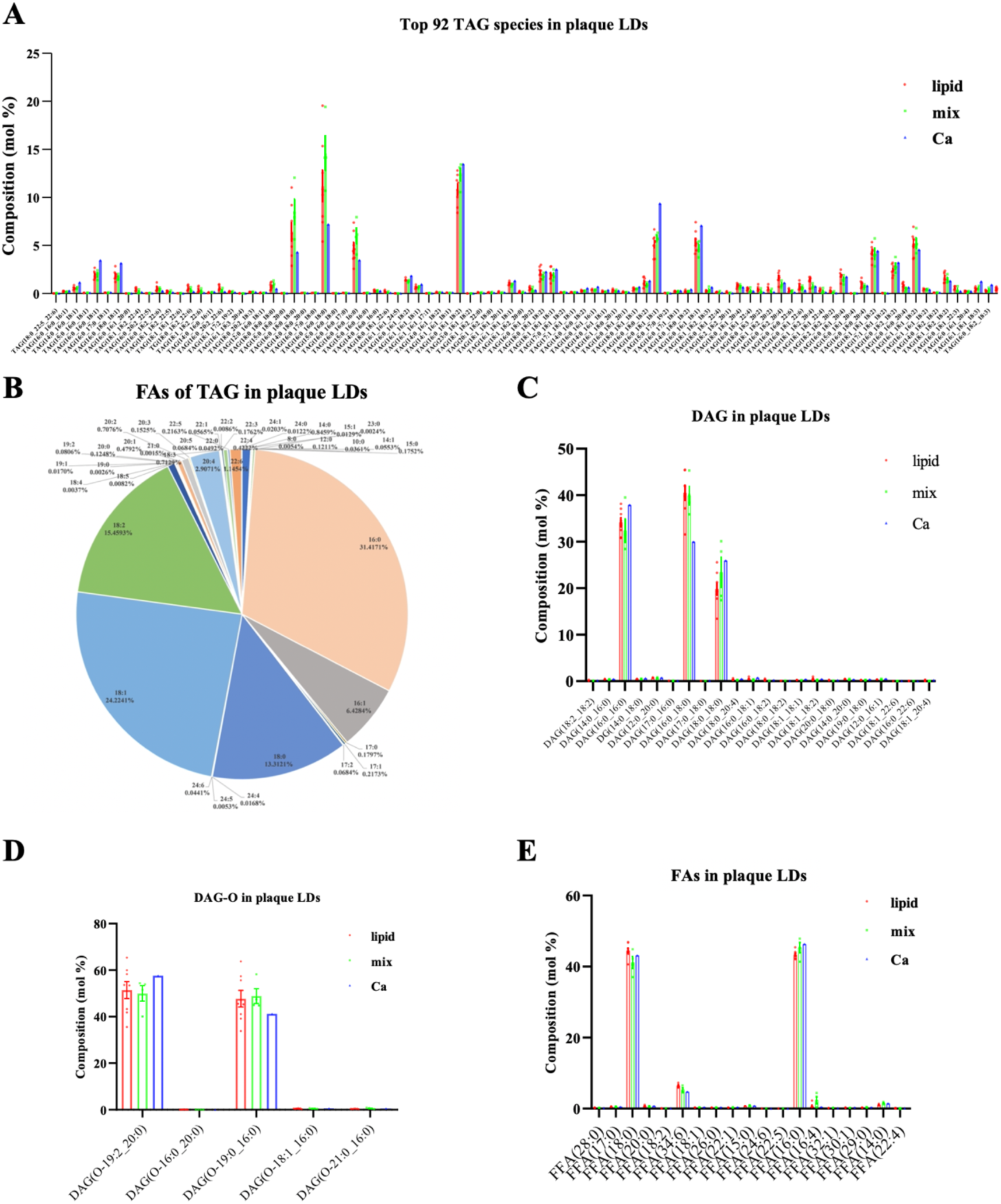
Different neutral lipid compositions of lipid droplets in three types of carotid artery plaques. A. Composition of triacylglycerol (TAG) species in LDs of three types of carotid artery plaques. B. Overall fatty acid composition of TAG in plaque LDs. The mole percentages of different fatty acid composition of TAG represent the average of a total of 13 LD sample. C-E. Composition of diacylglycerol (DAG), ester bond-diacylglycerol (DAG-O), and free fatty acid (FFA) species in LDs of three types of carotid artery plaques.

**Figure S4.**
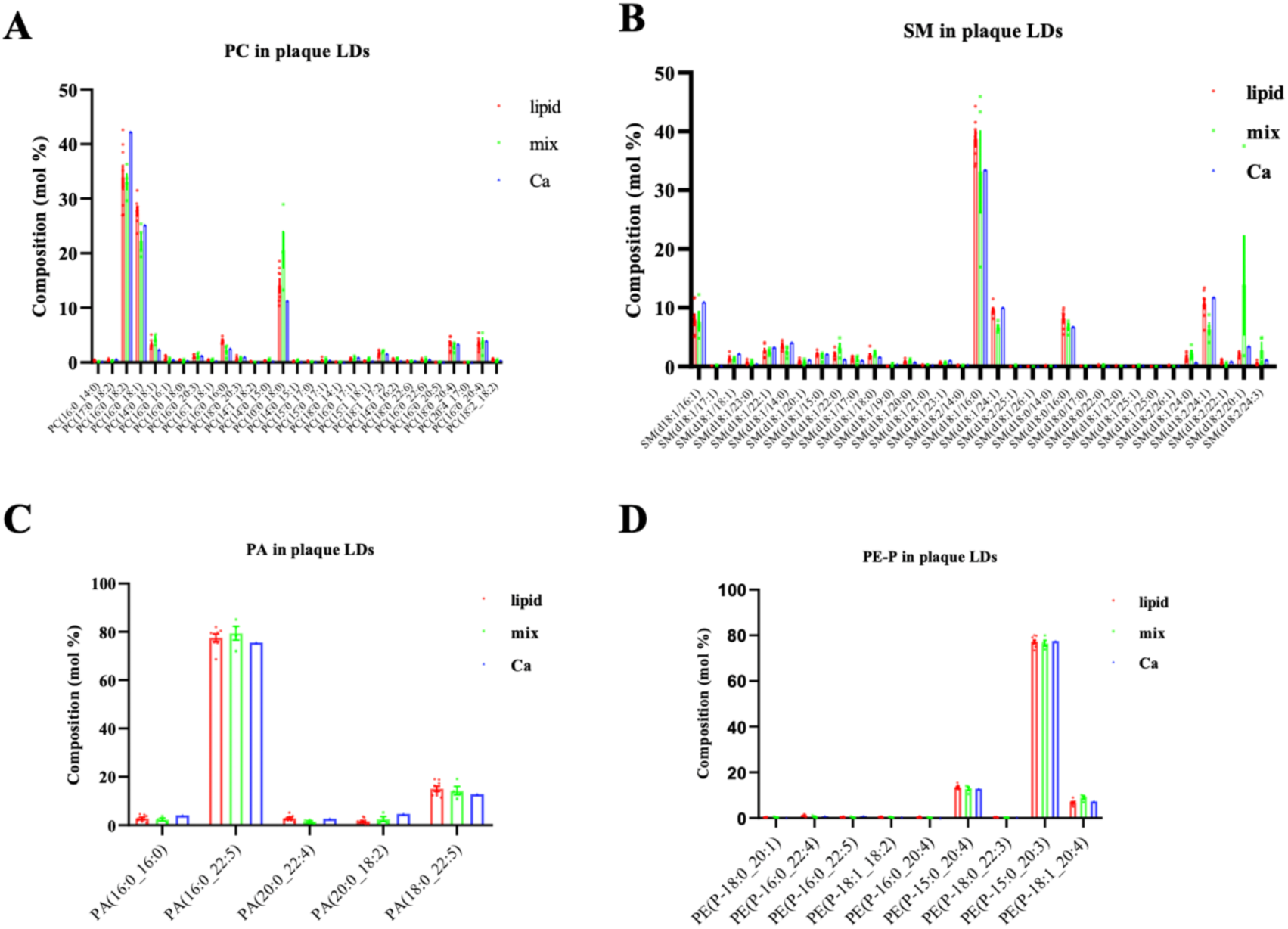
Different phospholipid compositions of lipid droplets in three types of carotid artery plaques. A-D. Composition of phosphatidylcholine (PC), sphingomyelin (SM), phosphatidic acid (PA), phosphatidylethanolamine (PE) and Plasmalogens (PE-P) species in LDs of three types of carotid artery plaques.

## Notes

### Competing Interest Statement

The authors have declared no competing interest.

